# Anti-HVEM mAb therapy improves antitumoral immunity both *in vitro* and *in vivo*, in a novel transgenic mouse model expressing human HVEM and BTLA molecules challenged with HVEM expressing tumors

**DOI:** 10.1101/2022.11.04.515180

**Authors:** C. Demerle, L. Gorvel, M. Mello, S. Pastor, C. Degos, A. Zarubica, F. Angelis, F. Fiore, J.A. Nunes, B. Malissen, L. Greillier, G. Guittard, H. Luche, F. Barlesi, D. Olive

## Abstract

**Background:** TNFRF-14/HVEM is the ligand for BTLA and CD160 negative immune co-signaling molecules as well as viral proteins. Its expression is dysregulated with an overexpression in tumors and a connection with tumors of adverse prognosis.

**Methods:** We developed C57BL/6 mouse models co-expressing human huBTLA and huHVEM as well as antagonistic monoclonal antibodies (mAbs) that completely prevent the interactions of HVEM with its ligands.

**Results:** Here, we show that the anti-HVEM18-10 mAb increases primary human αβ-T cells activity alone (CIS-activity) or in the presence of HVEM-expressing lung or colorectal cancer cells *in vitro* (TRANS-activity). Anti-HVEM18-10 synergizes with anti-PD-L1 mAb to activate T cells in the presence of PDL-1 positive tumors, but is sufficient to trigger T cell activation in the presence of PD-L1 negative cells. In order to better understand HVEM18-10 effect *in vivo* and especially disentangle its CIS and TRANS effects, we developed a knock-in (KI) mouse model expressing human BTLA (huBTLA^+/+^) and a KI mouse model expressing both human BTLA and human HVEM (huBTLA^+/+^ /huHVEM^+/+^ (DKI)). *In vivo* pre-clinical experiments performed in both mouse models showed that HVEM18-10 treatment was efficient to decrease human HVEM+ tumor growth. In the DKI model, anti-HVEM 18-10 treatment induces a decrease of exhausted CD8^+^ T cells and regulatory T cells and an increase of Effector memory CD4^+^ T cells within the tumor. Interestingly, mice which completely rejected tumors (± 20%) did not develop tumors upon re-challenge in both settings, therefore showing a marked T cell-memory phenotype effect.

**Conclusions:** Altogether, our preclinical models validate anti-HVEM18-10 as a promising therapeutic antibody to use in clinics as a monotherapy or in combination with existing immunotherapies (anti-PD1/anti-PDL-1/anti-CTLA-4).

## Introduction

Immunotherapies (IT) and immune checkpoint inhibitors (ICI) opened a new era in cancer treatment. ICI unleash anti-tumoral immune responses, leading to major clinical benefits in subgroups of patients. However, rate and duration of IT successes are variable among cancer types and patients. Indeed, in lung cancer anti PD-1/PD-L1 antagonistic antibodies offer a durable remission in 30% of patients (1,2). However, in colorectal cancer, the efficacy of anti-PD1/PD-L1 was restricted to a small subset of patients presenting a high microsatellite instability (MSI) status (3,4). Consequently, new IT targets and approaches are needed to achieve efficacy in a larger proportion of patients.

HVEM, or TNFRSF14, is a TNF-receptor family member largely expressed by healthy immune and non-immune cells and participates to immune homeostasis (5,6). HVEM is expressed on immune cells and also upregulated in numerous solid and hematologic malignancies such as melanoma (7), digestive cancers (8,9) or breast cancer (10). HVEM network of interactors is complex, and induces either cell activation or inhibition (11). Indeed, HVEM binding to BTLA (B and T lymphocyte attenuator) and CD160 (BY55) triggers co-inhibitory signals whereas its binding to LIGHT (TNFSF14) and Lymphotoxin-α (LTα) are co-stimulatory ligands. Similar to PD-1 and CTLA-4, BTLA is an important co-inhibitory receptor expressed by B and T cells (12). Therefore, targeting HVEM is a promising but complex IT strategy (13).

In our study, we focused on two of the most worldwide common cancers in both men and women: lung and colorectal cancer. In colorectal cancer, HVEM upregulation in malignant lesions is linked to tumor status and pathological stage, with an independent prognostic value (8). In lung cancer, HVEM expression seems to be a tumor driven mechanism, independent from the PD-L1 network that may contribute to immune escape (14).

Our team previously described a monoclonal anti-HVEM mAb (anti-HVEM18-10) that preferentially inhibits HVEM interaction with BTLA and enhances γδ-T cells responses against lymphoma (15). Here, we further evaluate the activity of anti-HVEM18-10 on immune cells and its anti-tumoral effect in pre-clinical T cell activity alone (CIS-activity) or in the presence of HVEM-expressing lung or colorectal cancer cells *in vitro* (TRANS-activity).

We also show the interest of antiHVEM18-10 combination with other immune checkpoint inhibitors such as antiPD-L1 mAb, although aHVEM18-10 is sufficient to trigger T cell activation in the absence of PDL1 expression. In order to better understand HVEM18-10 mAb effect *in vivo we* developed an innovative syngeneic immunocompetent mouse models expressing human BTLA (huBTLA^+/+^) or both human BTLA and human HVEM (huBTLA^+/+^huHVEM^+/+^, DKI). Our experiments performed in both mouse models showed that anti-HVEM18-10 treatment was efficient to decrease tumor growth, and strengthened local immune response. Moreover, the re-challenge of tumor-free mice (20% of treated mice) demonstrate that the immune response is durable with marked memory T cells phenotype. Altogether, our preclinical data validate anti-HVEM18-10 as a promising antibody to use in clinics alone or in combination with existing therapies (anti-PD1/anti-PD-L1/anti-CTLA-4).

## Methods

### Peripheral Blood Mononuclear Cells (PBMCs)

Human peripheral blood was obtained from Etablissement Francais du Sang (EFS) after obtaining informed consent from the donor. Human peripheral blood mononuclear cells (PBMC) were isolated using a Ficoll media (Eurobio) centrifuged at 800 × g for 30 min at room temperature with no break or acceleration. Cells were recovered from the interface with plasma, washed twice in PBS, counted and frozen in RPMI supplemented with 20% FCS and 10% DMSO until experimentation.

### Tumor Cell Lines

NCIH2291 and NICH2405 are lung adenocarcinoma cell lines purchased from ATCC and grown in RPMI supplemented with 10% FCS. HT29 is a colorectal cancer cell line grown in DMEM supplemented with 10% FCS. All cell lines were confirmed to be free of mycoplasmas using the MycoAlert detection kit (Lonza, Basel, Switzerland). Cells were detached in PBS EDTA 5mM without enzymatic solution to avoid HVEM cleavage.

### Antibodies

Anti-HVEM 18.10 and anti PD-L1 3.1 are homemade antibodies that have been described previously (15),(16). Briefly, mAbs are murine IgG1 anti-human HVEM or PD-L1 mAb, produced as ascites and purified by protein A binding and elution with the Affi-gel Protein A MAPS II Kit (Bio-Rad, Marnes-La-Coquette, France). Mouse IgG1 isotype control was purchased from Miltenyi Biotec (Bergisch Gladbach, Germany).

### HVEM and PD-L1 expression

Phenotypic expression was assessed by flow cytometry. HVEM expression was measured on CD4^+^ and CD8^+^ T cells in resting or activated conditions. 100 000 PBMC per well were distributed in 96 wells flat bottom plate, with donor specific OKT3 concentration (Ultra-LEAF Purified anti-human CD3 Antibody, Biolegend) and the following mABs when indicated: IgG1, anti-HVEM18-10, and/or anti-PD-L1 3.1 at 10 μg/ml in a final volume of 200μL RPMI supplemented with 10% FCS and 30 UI/ml IL-2 (Proleukine, Novartis). Noteworthy, OKT3 concentration was determined in advance for every PBMC donor, ranging from 5 to 50 pg/ml to obtain a sub-optimal T cells activation. Negative controls were PBMC without OKT3 and mAbs. T cell differentiation subsets were determined as: naive (CD27^+^CD45RA^+^), effector memory (CD27^+^CD45RA^-^), central memory (CD27^-^CD45RA^-^) and TEMRA (CD27^-^CD45RA^+^) (Table1). HVEM and PD-L1 expression on tumor cell lines was established twice with appropriate isotype control and viability staining.

**Table 1:** Flow Cytometry Panel for HVEM expression on T cells subsets.

| Marker | fluorochrome | Isotype |
| --- | --- | --- |
| CD27 | BV605 |  |
| CD45RA | BV711 |  |
| CD45 | BV786 |  |
| CD8 | PE |  |
| 7AAD | PeCy5 |  |
| CD56 | PeCy7 |  |
| HVEM | APC | IgG1-APC |
| CD3 | APCCy7 |  |

### Transcriptomic expression of HVEM and PD-L1

To study the transcriptomic expression of HVEM and PD-L1 data were extracted from The Cancer Genome Atlas (TCGA) database containing 526 lung adenocarcinoma samples and 388 colorectal carcinoma samples. Tumor cell lines gene expression values were extracted from the Cancer Cell Line Encyclopedia (CCLE). Data were normalized and expressed as log2(TPM+1) of raw value.

### Co-culture and proliferation assays

After thawing and overnight resting in RPMI supplemented with 10% FCS, PBMC were stained with Cell Trace Violet (CTV, Thermofisher) according to manufacturer instructions. Briefly PBMC were washed in PBS twice. CTV staining was performed at 37°c and 5% CO_2_ for 15 minutes precisely with 1μL CTV for 10 to 15 million PBMC/ml of PBS. Then, PBMC were washed twice in RPMI supplemented with 10% FCS. PBMCs were stimulated as described above. Briefly, PBMC plated and the following mABs: IgG1, anti-HVEM18-10, and/or anti-PD-L1 3.1 and IL-2. When tumor cell lines were cultured with PBMC, they were seeded in wells 24 hours before experimentation with 50 000 cells per wells to obtain a confluence around 60%. NCIH2405 and HT29 were treated with mitomycin C at 10μg/ml (Sigma Aldrich) during 3 hours and washed three times before adding PBMC. After 72H incubation (37°c, 5% CO2) PBMC were recovered, washed in PBS and stained at 4°c during 20 minutes with viability staining and the following antibodies: CD45, CD3, CD4, CD8, TCR-γδ, CD25, (Table 2). Acquisition was performed on FACS LSR II (Becton Dickinson). Application settings and sphero-beads (BD Biosciences) were used to ensure reproducible results between experiments. Data acquisition on LSRII was performed with BD DIVA software and data analysis was conducted with FlowJo (Treestar, Becton-Dickinson) software v.10.

**Table 2:** Flow Cytometry Panel for Proliferation Assay.

| Marker | fluorochrome |
| --- | --- |
| CD45 | BV786 |
| CD3 | PeCF594 |
| CD4 | BV605 |
| CD8 | APC |
| CD56 | PeCy7 |
| Tgd | PE |
| 7AAD | PeCy5 |
| CD25 | FITC |
| CTV | V450 |

**Table 3:** Panel for TILs exploration with Mass cytometry (Ext238)

| Marker | Metal |
| --- | --- |
| CD11b | 115In |
| CD4 | 141Pr |
| CD357 | 143Nd |
| CD45R | 144Nd |
| CD69 | 145Nd |
| CD8a | 146Nd |
| CD62L | 147Sm |
| CD278 | 148Nd |
| CD5 | 149Sm |
| Ly6C | 150Nd |
| CD25 | 151Eu |
| CD3e | 152Sm |
| CD366 | 153Eu |
| CD152 | 154Sm |
| CD317 | 155Gd |
| CD223 | 156Gd |
| FoxP3 | 158Gd |
| Rorgt | 159Tb |
| Tbet | 160Gd |
| CD279 | 161Dy |
| KI-67 | 162Dy |
| Klrg1 | 163Dy |
| TIGIT | 164Dy |
| MERTK | 165Ho |
| CD26 | 166Er |
| F4/80 | 167Er |
| XCR1 | 168Er |
| TCRβ | 169Tm |
| CD161 | 170Er |
| CD44 | 171Yb |
| Ly-6G | 172Yb |
| CD172a | 173Yb |
| MHCII | 174Yb |
| NKG2a | 175Lu |
| CD134 | 176Yb |
| CD45.2 | 198Pt |
| Cd11c | 209Bi |

**Table 4:**
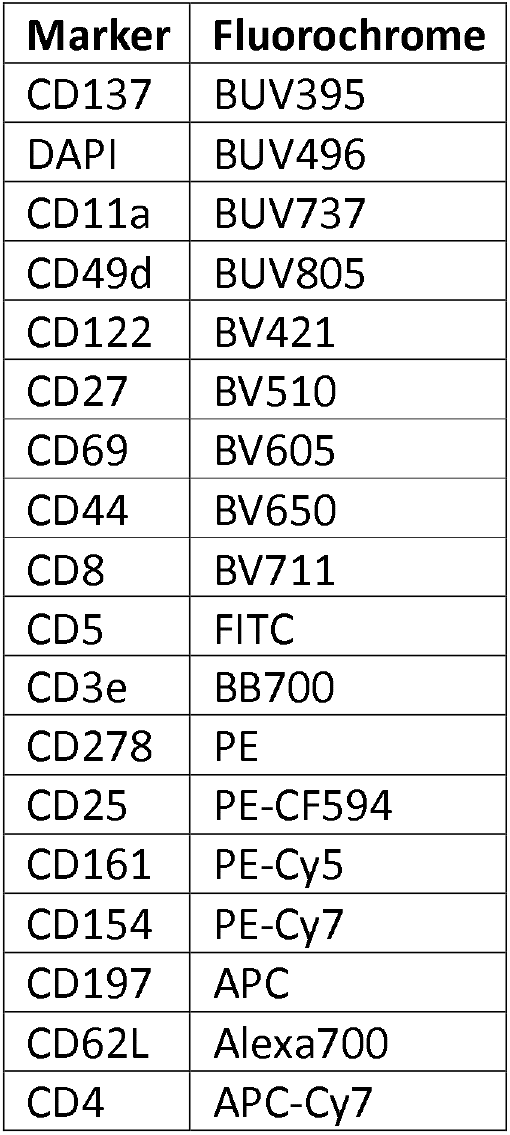
Flow cvtometry panel for lymph node phenotvping.

### Knock-in mouse model

To produce B6-Tnfrsf14^tm1Ciphe^ KI mice in which the mouse *tnfrsf14* exon 1 was replaced with the human *tnfrsf14* exon 1, ES cells were electroporated with the vector 1477_hTNFRSF14_03_v1 (containing the human cDNA sequence TNFRSF14-001 ENST00000355716 corresponding to human TNFRSF14 exon 1). ES cells were then cultured 8 days in 96 well plate under G418 selecting conditions (250μg/mL) and screened for proper vector integration by PCRs using primers 1477_ScES_RH5_Fwd: CTCCACTGCTGCTGCTGCTCTT and 1477_ScES_RH5_Rev: GTCCCCAAACTCACCCTGAA. Southern blot analysis was used to confirm the unique integration to the correct locus. ES validated clones were then injected into Balb/cN blastocysts. Germ-line transmission, proper deletion of the Neo cassette and the presence of the sequence coding for human Tnfrsf14 was assessed by PCR (see genotyping). To produce Btla^TmlCiphe^ mice, exon 2 from WT mouse was substituted by human exon 2.(17) The same method as described above was used to replace murine exon 2 with Btla-001 (ENSMUST00000102802) sequence. B6-Tnfrsf14^tm1Ciphe^ and Btla^Tm1Ciphe^ mice were then crossed to obtain double KI mice.

### Genotyping

HuBTLA^+/+^HuBTLA^+/+^ genotyping was performed using the following PCR primers: Fwd, 5’- TGCAATGATACCTATGGTCC −3’; Rev, 5’- TGACTGTTCTGATCTGGGG −3’; with expected band sizes at 537 bp for WT alleles and 616 bp for KI. HVEM ^hu KI^ genotyping was performed using the following PCR primers: Fwd KI, 5’- CCTTACATGTTTTACTAGCCAG −3’; Fwd WT, 5’- CTGCCTCTAACAGACTTCAGT −3’; Rev, 5’- TGAAGGTGTTGTCTGTAGGG −3’; with expected band sizes at 198 bp for WT alleles and 259 for KI.

### Experimental tumor experiments

Single-cell suspensions of MC-38 or MC-38^hu HVEM^ colorectal cancer cells were injected (0.5×10^6^ for neo challenge, 2×10^6^ for re-challenge) subcutaneously in the right flank of huBTLA^+/+^huBTLA^+/+^ mice. Mice bearing tumor between 50-100 mm^3^ where then randomized at day 9 and mice were injected with an isotype control (2mg/kg or 10mg/kg) or anti-HVEM18-10 (2mg/kg or 10mg/kg) or CTLA-4 antibody (2mg/kg) every 3-4 days with for a total of 6 injections.

### Mass cytometry for mice immunoprofiling

Tumors were resected from huBTLA^+/+^ and DKI mice which received anti-HVEM18-10, anti-CTLA-4 or isotype treatment. Tumors were collected and digested using the Tumor Dissociation Kit (Miltenyi Biotech). Digested tumors were mechanically disrupted using the GentleMACS Octo Dissociator (Miltenyi Biotech) to obtain a single-cell suspension, followed by isolation of CD45+ TILs using the CD45 (TIL) MicroBeads kit (Miltenyi Biotech). The cells were counted before proceeding for cell surface staining. Then, tumor cells were stained for viability using Cisplatin for 10 min at 37°C, washed and stained using the panel described in supplemental table 3. Finally, cells were incubated in the presence of Iridium, a DNA intercalator allowing the identification of cells. Stained cells were then acquired on a Helios mass cytometer (Cy-TOF, Fluidigm) and analyzed using the OMIQ software platform (OMIQ).

### Unsupervised CyTOF data clustering and phenotypic analysis

CyTOF data files were exported (Helios program, Fluidigm), debarcoded and live cells were gated in FlowJo (Treestar, BD). Live cell *.fcs* files were exported and analyzed using OMIQ online platform (OMIQ) (18). T cells were manually identified (TCRb^+^CD3^+^). T cell-gated data were subsampled to maximum equal available cell number (6000 T cells) and were subjected to an arcsinh transformation (co-factor 5). Clustering or tumor infiltrating T cells was performed using PhenoGraph (19), with the following parameters: Euclidean distance metric with K (nearest neighbor factor) =30 for cluster identification at the per mouse level. PhenoGraph clusters (n=19) were gated and displayed on a UMAP (Euclidian distance, neighbor factor 15, minimum distance 0.4) for phenotypic analysis. T cell marker expression was represented in a heatmap in function of the 19 clusters, where marker expression (columns) and clusters (rows) were subjected to hierarchical clustering using Euclidian distance (OMIQ).

### Flow cytometry for draining lymph node phenotyping

Mice which completely rejected tumors after re-challenge had their draining lymph node (LN) resected. LNs were dissociated and stained using the panel in supplemental table 4 Stained cells were then acquired on a LSR II cytometer (Becton-Dickinson) and analyzed using the OMIQ software platform (OMIQ).

### Statistical analysis

GraphPad Prism software was used to analyze and graph samples. For multiple comparison, two-way ANOVAs were performed and for comparison of condition pairs mann-whitney test was performed. *: p-val<0.05; **: p-val<0.01; ***: p-val<0.005; ****: p-val<0.001.

## Results

### HVEM is highly expressed on T cells and the blockade of HVEM in CIS enhances T cell activation

HVEM is known to be widely expressed among immune cells (6). Therefore, we studied the effect of anti-HVEM18-10 blockade directly on T cells (CIS blockade). PBMCs were isolated from healthy donors and stimulated for 72hrs with anti-CD3 in combination with anti-HVEM18-10 or an IgG1 control mAbs. We found that HVEM was highly expressed on activated CD4^+^ and CD8^+^ T cells (Fig1. A-B). HVEM expression is increased upon stimulation in CD4^+^ T cells (activated: 82.4 ± 3.1%; resting: 70,3 ± 5,7%) (Fig1. A), this effect was marked on CD4^+^ effector memory T cells (activated: 79.9 ± 4.9%; resting: 58.0 ± 5.6) (Fig1. C). HVEM expression did not differ on CD8^+^ T cell, upon stimulation or memory subtypes (Fig1. D). T cell proliferation and activation marked by the expression of CD25 significantly increased upon anti-HVEM stimulation compared to control (Fig.1 E-H). Taken together, our results show that CIS-HVEM blockade enhances T cell activation.

**Figure 1:**
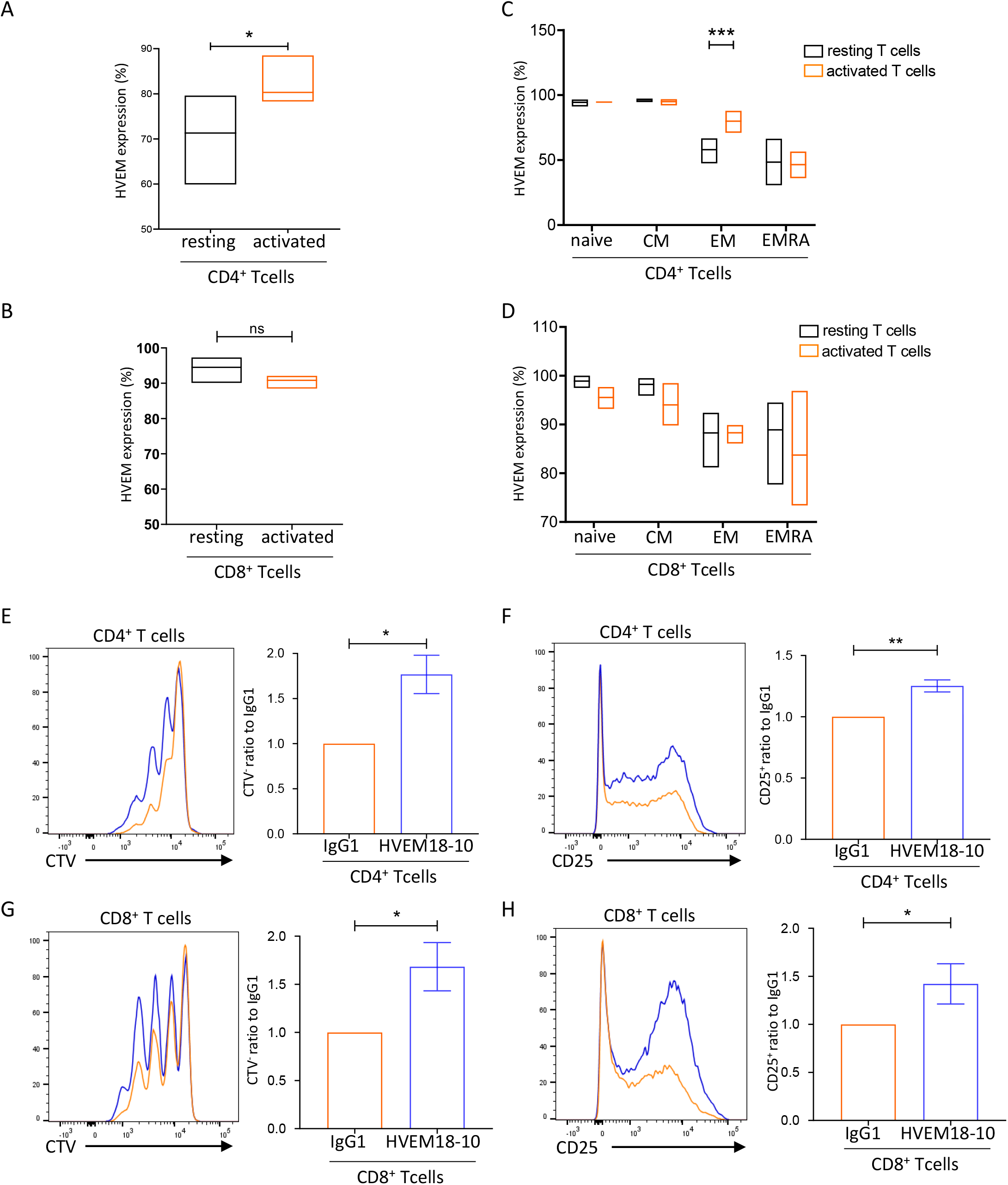
HVEM is highly expressed on T cells and the CIS-HVEM blockade enhance T cells activation. (A-H) PBMC were isolated from healthy donors, and cultured for 72h with anti-CD3 (OKT3) stimulation alone (orange bars), with OKT3 stimulation and anti-HVEM18-10 treatment (blue bars/lines), or without any stimulation (black bars). HVEM expression was assessed in resting (black bars) and activated (orange bars) CD4^+^ (A), CD4^+^ T cells subsets (naive, central memory, effector memory, and TEMRA) (B), CD8^+^ (C), CD8^+^ T cells subsets (D) by flow cytometry on healthy donors (n=3). (E-H) PBMC were incubated with OKT3 and anti-HVEM 18-10 antibody (blue bars/lines) or a control mAb IgG1 (orange bars/lines). proliferation profile of T cells was assessed by flow-cytometry detecting Cell Trace Violet (CTV) (E, G for CD4+ and CD8^+^ T cells, respectively) or CD25 staining (F, H for CD4^+^ and CD8^+^ T cells, respectively). (E, F, G, H) One representative plot is shown for each T cell subset. Bar plots are the Mean ± SEM of 4 different healthy donor samples. \**p*. val < 0.05, \*\**p*. val < 0.01, \*\*\**p*. val < 0.001 (Student’s t-test).

### HVEM expression is higher than PD-L1 in human lung and colorectal cancers

Next, we decided to study the expression status of HVEM on tumor from different cancers. Transcriptomic data from the TCGA containing 525 adenocarcinoma samples and 300 colorectal carcinoma samples were extracted, and the expression of HVEM and PD-L1 was examined (Fig2. A-B). We found that HVEM was highly expressed in both cancers. While PD-L1 expression was high in lung cancer and low in colorectal cancers. Interestingly, HVEM (lung: 17.69 ± 0.04; colorectal: 10.7 ± 0.03) expression is greater than that of PD-L1 (lung: 15.81 ± 0.06; colorectal: 4.2 ± 0.07) expression in both cancers (Fig2. A, B). Next, we studied HVEM and PD-L1 transcriptomic expression (from the cancer cell line encyclopedia) and cell surface expression by cytometry in lung and colorectal cancer cell lines (Fig2. C-D). HVEM and PD-L1 are heterogeneously expressed in lung cancer cell lines, we selected two lung cancer cell lines that expressed both genes (NCIH2291) or in large majority HVEM compared to PD-L1 (NCIH2405) for subsequent experiments (Fig2. C). The vast majority of colorectal cancer cell line highly expressed HVEM whereas PD-L1 was expressed in lower amounts (Fig2. D). Reflecting these observations, we selected HVEM^+^PD-L1^-^ HT29 cell line for subsequent experiments (Fig2. D). These results were confirmed in NCIH2291 (HVEM^+^ PD-L1^+^), NCIH2405 (HVEM^+^ PD-L1^-^) and HT29 (HVEM ^+^ PD-L1^-^) by flow cytometry assessment of PDL-1 and HVEM surface expression (Fig2. E, F).

**Figure 2:**
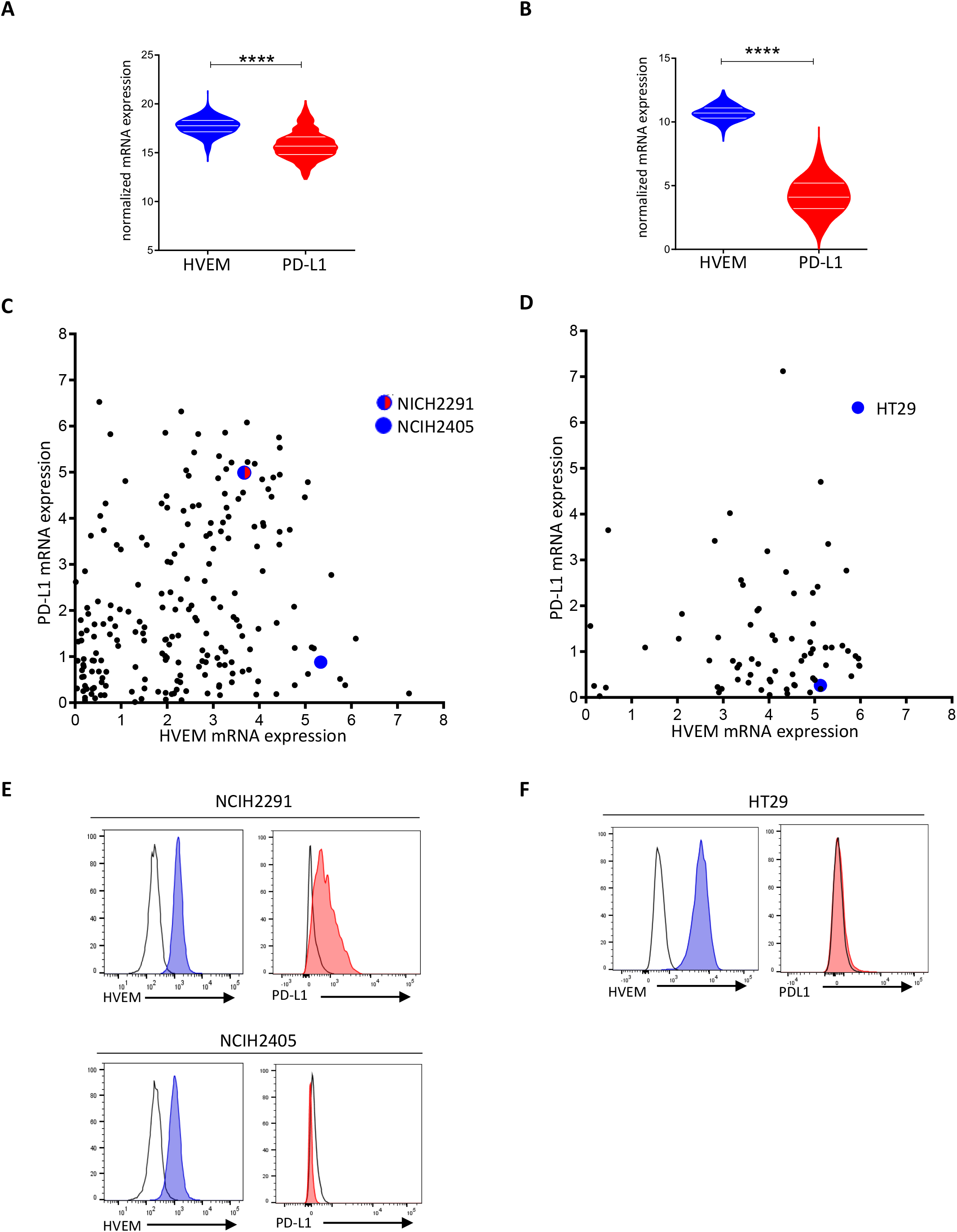
HVEM is more expressed than PD-L1 in lung and colorectal cancers. HVEM and PD-L1 transcriptomic expression was analyzed in 526 lung adenocarcinoma samples (A) and 388 colorectal carcinoma samples (D). Normalized expression data were extracted from The Cancer Genome Atlas (TCGA) database (**** *p*. val < 0,0001). HVEM and PD-L1 transcriptomic expression was analyzed in lung (B) and colorectal (E) cancer cell lines. (B, E) Data are extracted from the Cancer Cell Line Encyclopedia (CCLE) and expressed in log2(TPM+1). HVEM and PD-L1 phenotypic expression on selected Lung (C) (NCIH2291 and NCIH2405 respectively) and Colorectal (F) cancer cell line (HT29). HVEM staining in dark gray and control isotype in light gray.

### Anti-HVEM 18-10 enhances T cells responses against lung cancer cell line NCIH2291

Knowing the expression of HVEM among tumor cell lines allowed us to further test anti-HVEM effect. Co-cultures of PBMCs and the lung cancer cell line NCIH2291 (HVEM^+^ PD-L1^+^) were performed during 72 hours in the presence of anti-CD3 in combination with an IgG1 control or anti-HVEM18-10. The addition of anti-HVEM18-10 to the co-culture drastically increased proliferation ratio of CD4^+^ (1.83 ± 0.18) and CD8^+^ (1.64 ± 0.08) T cells (Fig3 A, C) compared to control. In the presence of NCIH2291, the stimulation by anti-HVEM18-10 induced a greater CD8^+^ T cell proliferation (CIS+TRANS effects, Fig3.A,B) compared to that observed in absence of NCIH2291 in the same conditions (CIS effect, Fig1.E,G). Using the same experiment settings, we observed an increase in the frequency of CD25 expression on CD4^+^ and CD8^+^ T cells in the presence of anti-HVEM18-10 compared to control (CD25 expression ratio CD4^+^ T cells: 1.27 ± 0.05; CD8^+^ T cells: 1.38 ± 0.06) (Fig3. C, D). Moreover, the addition of anti-HVEM18-10 to the co-culture also drastically increased the amount of pro-cytotoxic cytokines TNF-α (ratio 2.3 ± 0.41) and IFN-γ (ratio 8.0 ± 2.6) as measured by ELISA (Fig3.E, F). These results confirm an effect of the blockade of HVEM on CIS-T cell activation and Trans-activation as well.

**Figure 3:**
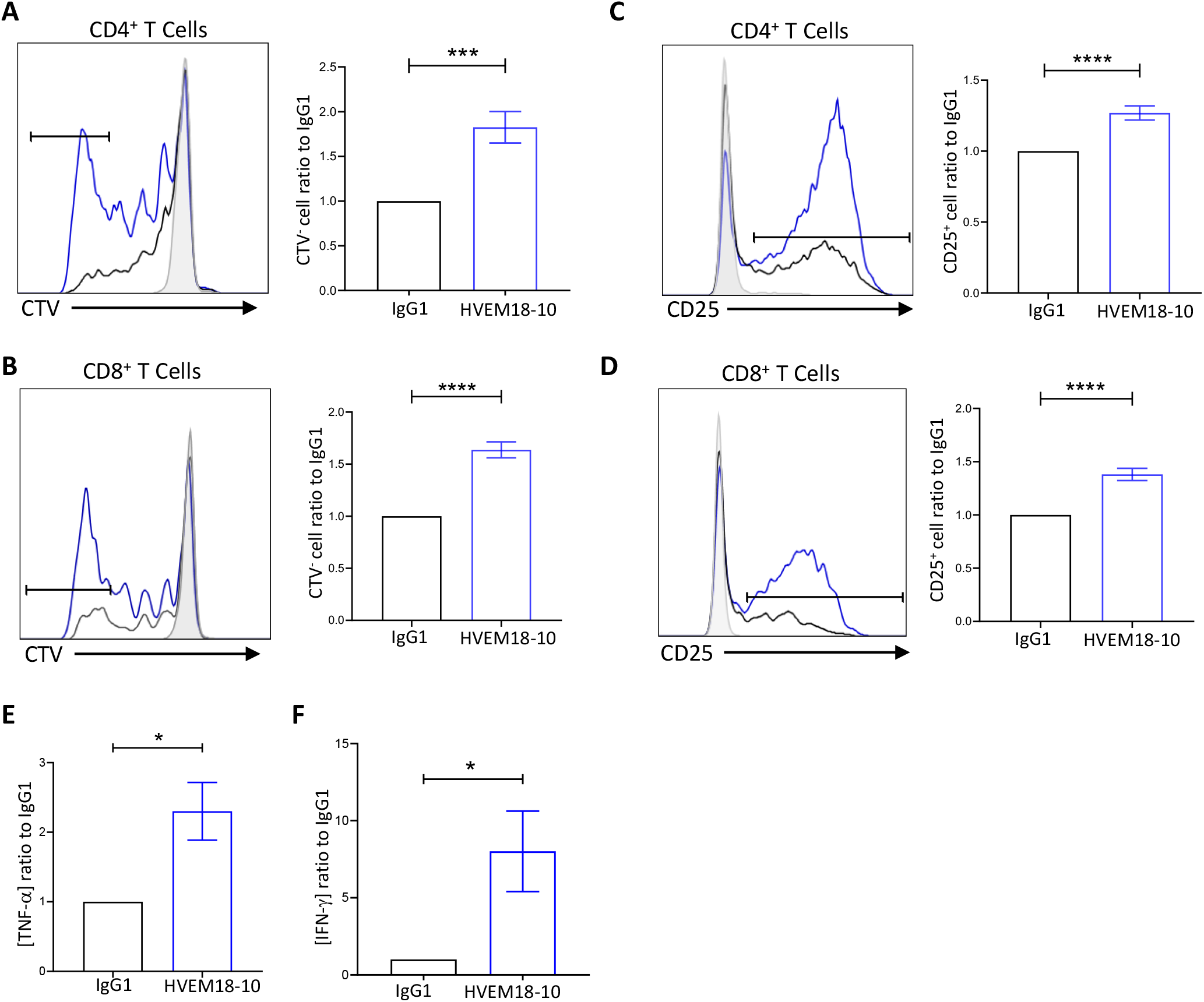
Anti-HVEM 18-10 enhances T cells responses against NCIH2291. Adherent lung cancer cell line NCIH2291 was seeded in wells 24hours before the experiment. Then we cultured PBMC from healthy donors for 72h with OKT3 stimulation and treated or not with anti-HVEM 18-10 antibody (blue bars/lines) (A, B) TNFα (A) and IFNγ (B) secretion was measured by Elisa in culture supernatant. (C-F) proliferation profile of T cells by Cell Trace Violet staining (C for CD4^+^ and D for CD8^+^ T cells) and CD25 expression (E for CD4^+^ and F for CD8^+^ T cells). Bar plots are the Mean ± SEM of different healthy donor samples, (A-B n=7; C-F n= 14). * *p*. val < 0.05, \*\**p*. val < 0.01, \*\*\**p*. val < 0.001, \*\*\*\**p*. val < 0.0001(Student’s t-test)

### Anti-HVEM18-10 triggers T cell activation, proliferation and synergizes with anti-PD-L1

Anti-PDL-1 antibody has been a real game changer in immunotherapy to treat lung cancer patients (20). Therefore, we thought to test whether the combination of anti-HVEM18-10 to anti-PDL-1 could improve T cell responses in co-culture experiments with HVEM and PD-L1 positive lung tumor cell line. PBMCs were primed with suboptimal anti-CD3 dose, then activated with anti-HVEM18-10, anti-PDL-1, both antibodies (combo), or IgG1 control during 72 hours (Fig 4A-D). The addition of anti-HVEM18-10 (Blue bar) was sufficient to improve T cells proliferation (CD4^+^ T cells proliferation ratio: 1.83 ± 0.18; CD8^+^ T cells: 1.64 ± 0.08) and CD25 expression compared to control (CD25 expression ratio CD4^+^ T cells: 1.27 ± 0.05; CD8^+^ T cells: 1.38 ± 0.06). Likewise, the anti-PDL-1 (Red bar) also increased T cell activation (CD25 expression ratio CD4^+^ T cells: 1.30 ± 0.08; CD8^+^ T cells: 1.54 ± 0.10) and proliferation ratio (CD4^+^ T cells: 1.95 ± 0.20; CD8^+^ T cells: 1.81 ± 0.09) compared to control. Interestingly, the combination of anti-HVEM18-10 and anti-PDL-1 (Purple bar) in culture showed even greater effect on T cell proliferation (proliferation ratio CD4^+^ T cells: 3.57 ± 0.21; CD8^+^ T cells: 3.06 ± 0.71) and CD25 expression (CD25 expression ratio CD4^+^ T cells: 1.60 ± 0.22; CD8^+^ T cells: 2.31 ± 0.20) compared to the separate conditions. These results highlight the potential of anti-HVEM18-10 and anti-PD-L1 combination to strengthen T cell activation.

**Figure 4:**
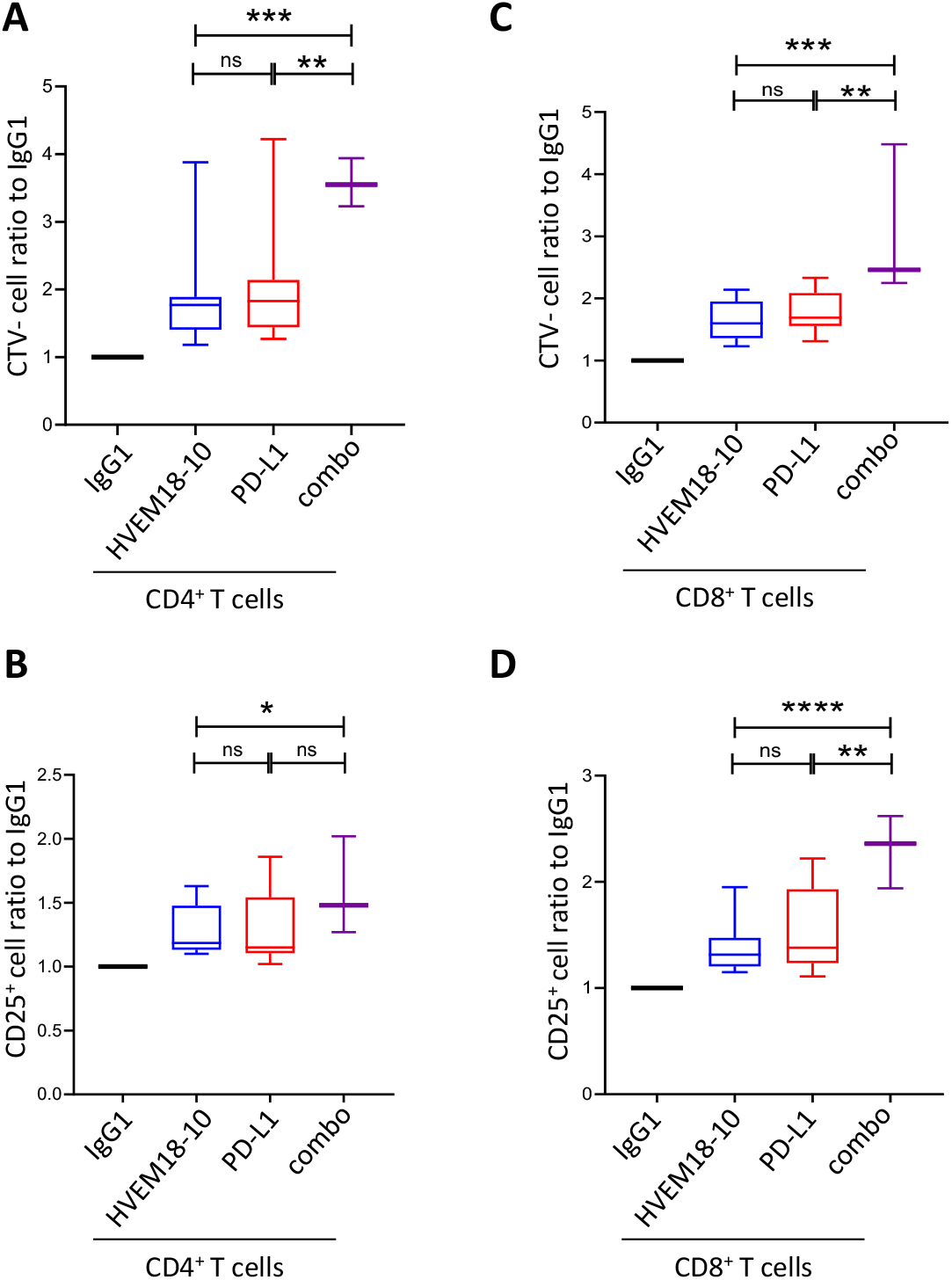
Anti-HVEM 18-10 synergizes with anti-PD-L1 to enhance T cells responses against lung cancer cell line NCIH2291. Adherent lung cancer cell line NCIH2291 were cultured with PBMC from healthy donors for 72h with CD3 stimulation and treated or not with anti-HVEM 18-10 antibody (blue bars/lines) (A-D) proliferation profile of T cells by Cell TraceViolet staining (A for CD4^+^ and B for CD8^+^ T cells) and CD25 expression (C for CD4+ and D for CD8+ T cells). (E,F) TNFα (E) and IFNγ (F) secretion was measured by Elisa in culture supernatant. Bar plots are the Mean ± SEM of different healthy donors samples, (A-D n= 14;E-F n=7). * p < 0.05, **p < 0.01, ***p < 0.001, ****p < 0-0001(Student’s t-test)

Interestingly, we next tested whether anti-HVEM18-10 could enhance T cell response against PD-L1^-^ lung and colorectal cancer cell line. Thus, we tested co-culture experiments with two HVEM^+^PD-L1^-^ cell lines from lung (NCIH2405) or colorectal cancer (HT29) in the same experiments setting as in Fig4. As expected, anti-PD-L1 treatment did not increase T cells activation in PD-L1^-^ lung nor colorectal co-cultures (Supp.Figure1). However, anti-HVEM18-10 was sufficient to improve CD4^+^ and CD8^+^ T cell activation with a marked increase of CD25 expression and proliferation in co-culture with either lung or colorectal cell line, validating the use of anti-HVEM18-10 with PD-L1^-^ tumors for future IT treatments. Taken together, our results demonstrate that not only anti-HVEM18-10 treatment induces T cell proliferation and activation *in vitro* in combination with anti-PD-L1, but represents a valuable alternative in PD-L1^-^ conditions.

### huBTLA^+/+^ and DKI Mice show WT-like hematopoietic cell proportions

To test whether the effect of anti-HVEM18-10 *in vivo* occurs via CIS- or TRANS-blockade using pre-clinical tumor models, we developed KI mice model expressing either human BTLA protein (huBTLA^+/+^) or human BTLA protein and human HVEM proteins (huBTLA^+/+^/huHVEM^+/+^or DKI) (Fig5. A). To produce huBTLA^+/+^ mice, exon 2 from WT mouse was substituted by human exon 2. 15. Likewise, HVEM^+/+^mice were developed by replacing exon 1 with human HVEM gene (tnfrsf14-001) exon 1. These mice were then crossed to obtain double KI (DKI) mice. The presence of the insertion was confirmed by PCR and cytometry. The expression of human/mouse HVEM on T cells, DC, Ly6C^+^cells and neutrophil and BTLA on T cells and B cells was examined by cytometry (suppl.Fig2 A,B). Both HuBTLA^+/+^ and DKI mice showed the same number of cells in the spleen (Fig 5B) and same proportion of T cells by cytometry. Other lymphoid and myeloid cells are also found in the same proportions in both models (Fig5.C). Indeed, the frequencies of B cells, γδ-Tcells and NK cells were similar among mice models. Similarly, myeloid cell frequencies (PMN, cDC1, cDC2, macrophages, monocytes) remained unchanged among these models (Fig5.C). Gating strategies for different cells subset are showed in suppl.Fig2 C. These observations allowed further *in vivo* tumor response studies with these new mouse models.

**Figure 5:**
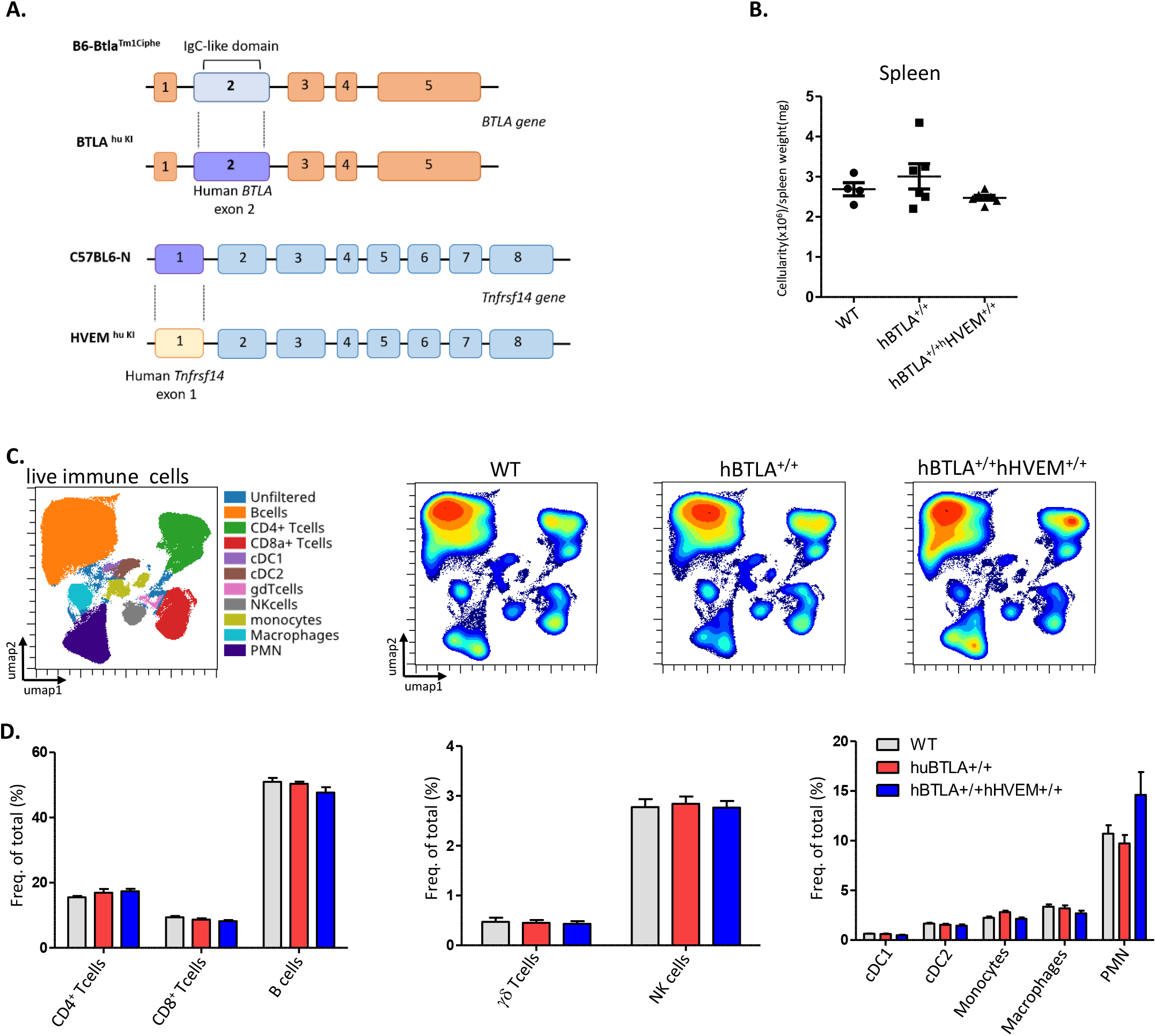
Hu BTLA and DKI Mice show the same hematopoietics cell homeostasis. A. representative scheme of huBTLA^+/+^huBTLA^+/+^ and huHVEM^+/+^ genetic modification. For huBTLA^+/+^huBTLA^+/+^ murine exon 2 was replaced by human BTLA exon2 and for huHVEM^+/+^murine exon 1 was replaced by human HVEM gene exon1. B. Numbers of total cell per spleen weight for WT, huBTLA^+/+^huBTLA^+/+^ and huHVEM^+/+^ is represented. C. UMAP showing the distribution of immune cell subset within WT, Hu BTLA and DKI mice (top left). D. Immune cell subset frequency was quantified in the 3 mice groups (top right). Density UMAP representing major and minor cell subsets (bottom).

### Blocking Trans-BTLA-HVEM binding *in vivo* is sufficient to decrease solid tumor growth

We then decided to challenge huBTLA^+/+^ mice with a colorectal cancer cell line MC-38 that is not expressing constitutively HVEM. 0.5 10^6^ MC-38 tumor cells where injected subcutaneously in the right flank of huBTLA^+/+^ mice (suppl.Fig3A). Mice bearing tumor between 50-100 mm^3^ were then randomized at day 9 and mice were injected with anti-HVEM18-10 or isotype control every 3-4 days with for a total of 6 injections (Fig6.A). As envisioned, tumor growth of parental MC-38 (HVEM) injected with an isotype control or with anti-HVEM18-10 at 10mg/kg did not show any significant decrease in tumor growth (Fig.suppl. 3B). We then transduced human HVEM in MC-38 cells and selected a clone that that was able to bind efficient anti-HVEM18-10 (namely: MC-38^hu HVEM^;suppl.Fig3A). As previously described, 0.5 10^6^ MC-38^hu HVEM^ tumor cells where injected subcutaneously in the right flank of huBTLA^+/+^ mice. Mice bearing tumor between 50-100 mm^3^ were then randomized at day 9 and mice were injected with an isotype control or anti-HVEM18-10 every 3-4 days with for a total of 6 injections. Here, we observed a decrease in tumor growth at any dose of anti-HVEM18-10 used (2 and 10 mg/kg) (Fig.6, B; Supp. Fig.3C). Moreover, 2 mice out of 10 in the 2 mg/kg dose and 1 out of 10 in the 10 mg/kg dose fully rejected the tumors. These 3 tumor free mice were re-challenged with 4-fold more concentrated MC-38^hu HVEM^ tumors cell inoculate (i.e. 2.10^6^ cells) in contralateral flank. All animals were monitored during 21 days post challenge to ensure immunological memory persistence. In 2 of these 3 mice the tumor was not measurable at day 21. However, after the euthanasia of those three mice at day 21, tumor mass was absent in their left flank revealing the presence of a scar tissue instead (Fig.6.C). Altogether these data confirmed the *in vivo* efficiency of anti-HVEM18-10 to block Trans-BTLA-HVEM binding.

**Figure 6:**
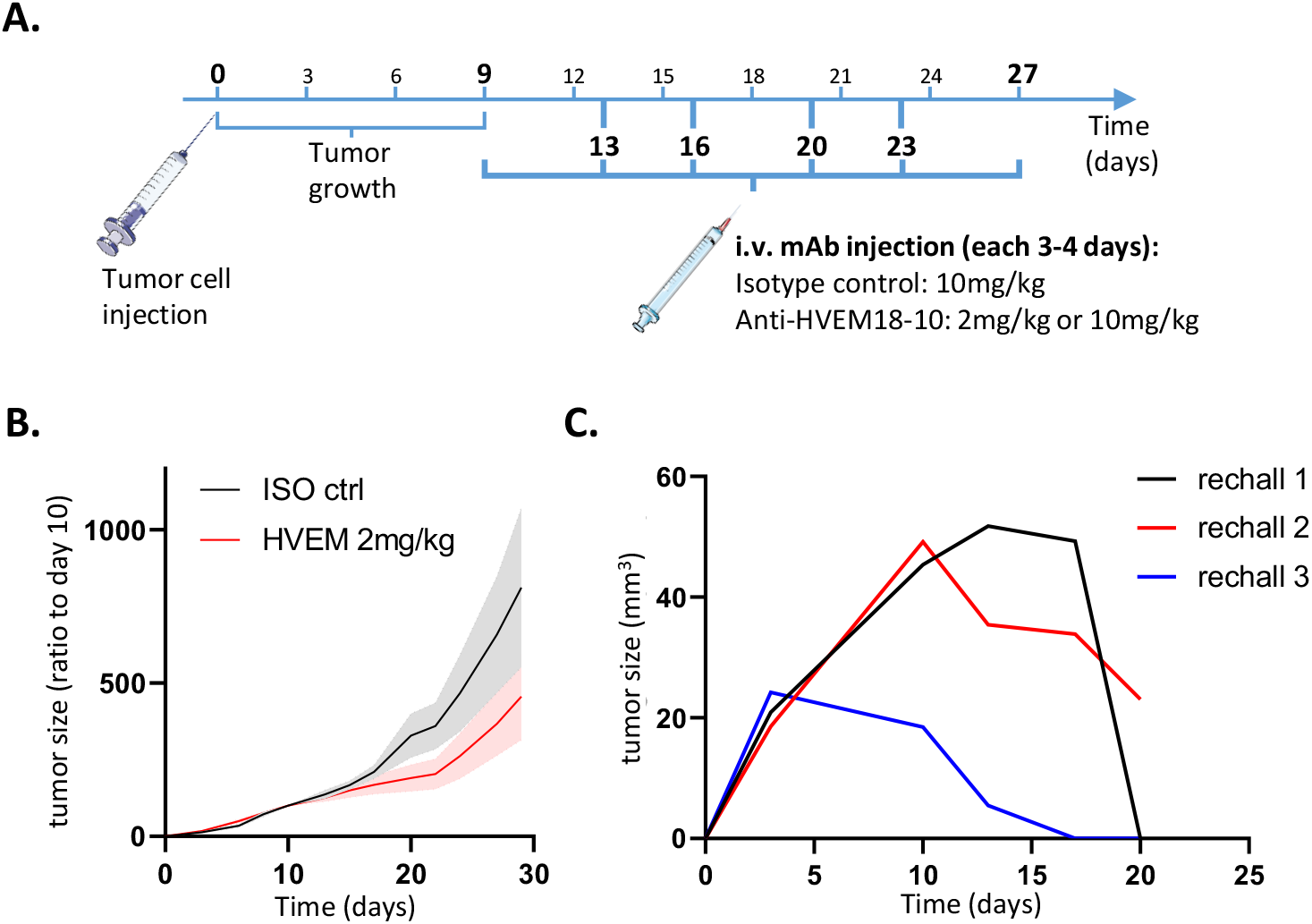
blocking Trans-BTLA-HVEM binding *in vivo* is sufficient to decrease solid tumor growth. A. Scheme representing *in vivo* tumor experiments settings. B. Measure of tumor growth showed as ratio to size of tumor at randomization day. Colorectal cancer cells MC-38^hu HVEM^ were injected (0.5×10^6^) at day 0, and Iso Ctrl (black) or anti-HVEM 18-10 antibody at 2mg/kg (red) concentration were injected. C. Tumor free mice from B and C measure of tumor growth after MC-38^huHVEM^ tumor cells re-challenge (2×10^6^). (B-C) n≥7; D n=3. \**p* < 0.05, \*\**p* < 0.01, \*\*\**p* < 0.001, \*\*\*\**p* < 0.0001 (two-way Anova).

### Anti-HVEM treatment decreases tumor growth, exhausted CD8^+^ T cell, KLRGl^-^ Treg infiltrate and increases EM conventional CD4^+^ T cell infiltrate

Then, we investigated the effect of anti-HVEM on lymphocytes to confirm whether DKI T cells were activated in a similar way than human T cells (Fig.7A). The proliferation of sorted spleen T cells was assessed after 72hrs of stimulations with coated anti-CD3 and anti-HVEM18-10 or isotype control at increasing concentration. The addition of anti-HVEM18-10 increased DKI T cells proliferation in a dose dependent manner similarly to human T cells (Fig.1), validating *in vitro* this new mouse model (Fig7.A). The effect of anti-HVEM18-10 *in vivo* was then studied using DKI mice. DKI mice were challenged with 0.5 10^6^ MC-38^hu HVEM^ tumor cells which were injected subcutaneously in the right flank. Mice bearing tumor between 50-100 mm^3^ were then randomized at day 6 or day 9 and mice were injected with an isotype control or anti-HVEM18-10 every 3-4 days for a total of 2 to 6 injections. We observed a decrease in tumor growth when anti-HVEM18-10 was injected at 2mg/kg (Fig7.B). Moreover, 3 mice out of 12 in the anti-HVEM18-10 completely rejected the tumors and were later investigated in re-challenge experiments. To understand the immune mechanisms underlying tumor reduction after anti-HVEM18-10 treatment, tumor infiltrating lymphocytes (TILs) phenotypes were investigated in anti-HVEM18-10 treated mice and isotype controls by mass cytometry. Tumor infiltrating T cells was mapped using UMAPs in anti-HVEM18-10 and isotype controls treated mice. Density analysis of UMAPs showed phenotypic variations between conditions (Fig7.C), that was investigated by unsupervised analysis using T cell subset markers (Supp.Fig4.A-C). Next, a clustering of tumor infiltrating T cells was performed using the well described PhenoGraph algorithm and the 19 identified cluster were projected on the UMAP highlighting the heterogeneity of intratrumoral T cells. 5 clusters belonging to Tregs (CD4^+^CD25^+^FoxP3^+^), 8 clusters belonging to conventional CD4^+^ T cells (conv. CD4^+^ T cells/non Tregs) and 6 clusters belonging to CD8^+^ T cells were identified (Fig7.D, E). Interestingly, while T cell clusters were very heterogeneous, only 3 clusters that where significantly altered were identified in anti-HVEM18-10 compared to isotype control treated mice (Fig7.E-H; Supp.Fig4B). These clusters were manually gated for validation in Supp.Fig4D,E). First, cluster 13, a subset of Tregs, was statistically decreased in anti-HVEM18-10 condition (Fig7.F; Supp.Fig4E). Cluster 13 was composed of KLRG1^-^CTLA-4^+^Ki67^+^ Tregs participating to the immunosuppression in the TME. Moreover, cluster 11, composed of effector memory (EM, CD44^+^CD62L^-^) CD4^+^ conv. T cells, was significantly increased under anti-HVEM18-10 treatment (Fig7G; Supp.Fig4E). In this cluster EM CD4^+^ T cells did not exhibit marks of immunosuppression or exhaustion such as high levels of PD-1, TIM-3 or TIGIT. Therefore, these EM CD4^+^ T cells may favor anti-tumoral immune response. Third, cluster 17, an EM (CD44^+^CD62L^-^) CD8^+^ T cell subsets showed higher expression of PD-1 and expressed TIM-3, LAG-3 and CTLA-4. Therefore, we labelled these EM CD8^+^ T cells as exhausted. Interestingly, these exhausted CD8^+^ T cells decreased under anti-HVEM18-10 treatment (Fig7.H; Supp.Fig4E). Taken together, our results demonstrate a loss in immunosuppressive T cell subsets (KLRG1^-^ Tregs, and exhausted EM CD8^+^ T cells) and an increase in EM CD4^+^ T cells. These phenotypic changes contribute to the reduction of tumor growth and correspond to phenotypes observed in previous studies showing tumor reduction following immunotherapeutic treatments in mouse models (21) (22).

**Figure 7:**
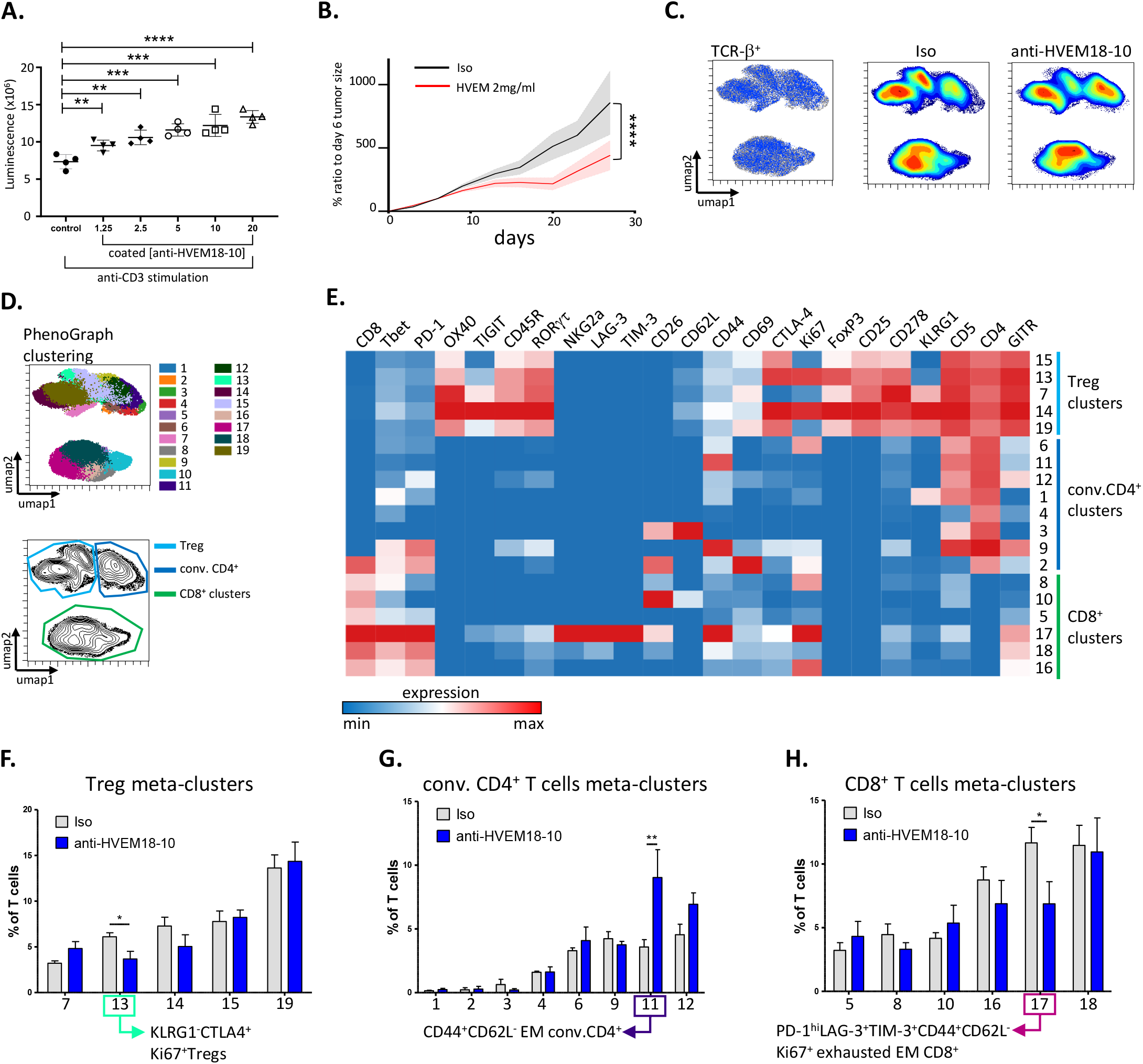
anti-HVEM treatment decreases tumor growth, exhausted CD8^+^ T cell, KLRGl^-^ Treg infiltrate and increases EM conventional CD4^+^ T cell infiltrate. A. T cells from DKI mice or WT were cultured with CD3 alone or with anti-HVEM 18-10 at increasing concentration. T cells proliferation assessed by luminescence. B. Measure of tumor growth showed as ratio to size of tumor at randomization day. Colorectal cancer cells MC-38^hu HVEM^ were injected (0.5×10^6^) at day 0, and Iso Ctrl (black) or anti-HVEM 18-10 antibody at 2mg/kg (blue) concentration were injected. C. Umap analysis of anti-HVEM18-10 (blue) or Isotype control (Iso, grey). Density analysis revealed phenotypic modification between anti-HVEM treatment and Isotype treatment. D. PhenoGraph clustering was performed on TILs from anti-HVEM18-10 and isotype control (iso) treated mice and represented as a Umap. 19 meta-clusters were identified among T cells (top). Treg (light blue), conventional CD4^+^ T cell (dark blue) and CD8^+^ T cell (green) cluster gating on a Umap (bottom). E. Heat-map representation of marker expression among the 19 identified PhenoGraph meta-clusters. Cluster were associated to Tregs (light blue), conventional CD4^+^ T cells (dark blue) and CD8^+^ T cells (green). Bar charts represent cluster variation between anti-HVEM18-10 treatment compared to Isotype control in Tregs (F.), conventional CD4^+^ T cells (G.) and CD8^+^ T cells (H.). \**p*.val <0.05; \*\**p*.val<0.01.

### Anti-HVEM-therapy builds a tumor specific memory T cell response associated with tumor antigen responsive T cells

We next decided to compare *in vivo* the effect of anti-HVEM response to a well-known checkpoint inhibitor anti-CTLA-4 on tumor growth and T cell activation. DKI mice were challenged with 2 10^6^ MC-38^hu HVEM^ tumor cells, which were injected subcutaneously in the right flank. Mice bearing tumor between 50-100 mm^3^ where then randomized at day 6 and mice were injected with an isotype control, anti-HVEM18-10 (2mg/kg) every or anti-CTLA-4 (2mg/kg) at day 7 and 10. Tumor infiltrating and draining lymph node T cell phenotype was analyzed on day 14. As expected, the injection of both anti-HVEM18-10 and anti-CTLA-4 decreased tumor growth (Fig8.A) and led to total tumor rejection in 3 mice under anti-HVEM treatment. Similarly, the injection of anti-CTLA-4 antibody led to total tumor rejection in 3 mice. Hence, tumor free mice respectively from anti-HVEM and anti-CTLA-4 treatment were re-challenged with 6.10^6^ cells MC-38^hu HVEM^ tumors cells in contralateral flank. Interestingly, tumors did not grow in these mice 14 days after re-challenge. Therefore, we hypothesized that primary tumor rejection built an anti-tumor T cell memory response that led to re-challenge rejection. Hence, we compared T cell response in re-challenged mice following anti-HVEM18-10 and anti-CTLA-4 treatments. Draining lymph nodes (LN) were resected and dissociated then T cell phenotypes were assessed by flow cytometry. T cell mapping was drastically modified after re-challenge compared to neo-challenge as shown by density UMAPs (fig8.B). Then, memory T cell subsets were gated (using CD44 and CD62L expression) and neo-challenged and re-challenged mice treated with anti-HVEM or anti-CTLA-4 were compared. We found that re-challenged treated mice either for anti-HVEM18-10 (23.86% ± 3.37) or anti-CTLA-4 (37.26% ± 8.7) were enriched in Effector Memory (EM) CD4^+^ T cells (CD44^+^CD62-L^+^) compared to their neo-challenged counterpart (anti-HVEM18-10: 10.29% ± 1.09; anti-CTLA-4: 12.96% ± 1.83) and isotype control (8.74% ± 1.07) (Fig8.C left). This increase was significantly superior in re-challenged CTLA-4-treated compared to rechallenged HVEM-treated mice (Fig8.C left). Noteworthy, naive CD4^+^ T cells decreased after re-challenge. Concerning CD8^+^ T cells, we found that re-challenged mice with anti-HVEM18-10 (48.2% ± 5.7) or anti-CTLA-4 (33.46% ± 7.07) showed more Central Memory (CM) CD8^+^ T cells (CD44^+^CD62-L^-^) compared to their neo-challenged counterpart (anti-HVEM18-10: 13.73% ± 1.38; anti-CTLA-4: 13.48% ± 0.88) and isotype control (16.38% ± 1.059) (Fig8.C left; Supp.Fig4D). This increase tended to be superior in re-challenged HVEM-treated compared to re-challenged CTLA-4-treated mice. Again, naïve CD8^+^ T cells decreased as well upon treatments. Taken together, these results show an enrichment in memory CD8^+^ and CD4^+^ T cells in re-challenged conditions.

**Figure 8:**
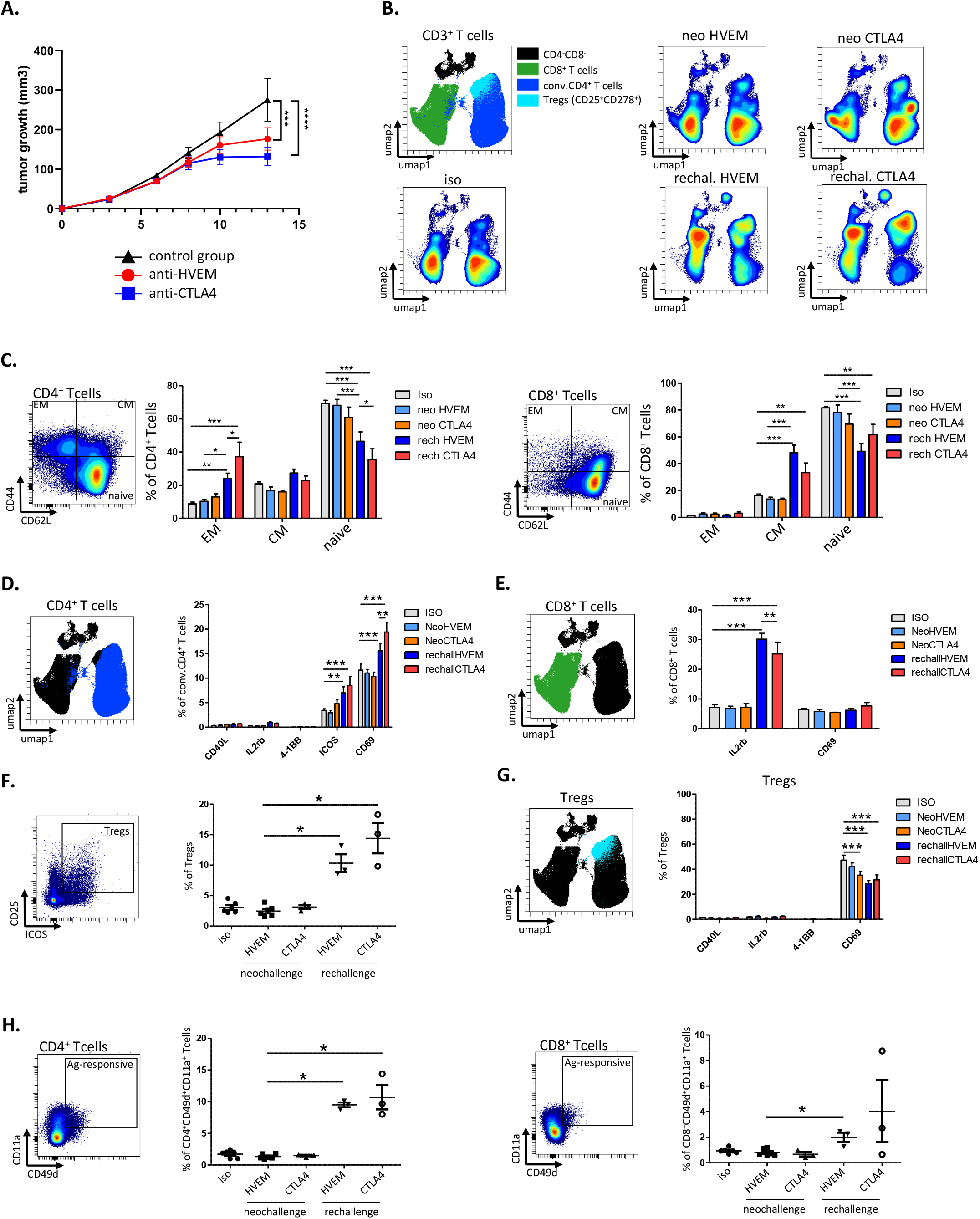
anti-HVEM18-10-therapy builds a memory T cell response associated with CD4^+^ and CD8^+^ tumor antigen responsive T cells. A. Tumor growth profile overtime following, isotype control (black), anti-CTLA-4 (blue) or anti-HVEM18-10 (red) treatment. B. lymphnodes (LN) were dissociated and cell stained for flow cytometry. UMAPs represent the phenotypic distribution of T cells within lymphnodes in isotype control mice, anti-HVEM18-10 or anti-CTLA-4 treated neo-challenged mice or anti-HVEM18-10 or anti-CTLA-4 treated re-challenged mice. C. Study of EM, CM and naive conventional CD4^+^ and CD8^+^ T cells among neo-challenged and re-challenged conditions D. Study of immune checkpoint expression in conventional CD4^+^ T cells among neo-challenged and re-challenged conditions. E. Study of immune checkpoint expression in CD8^+^ T cells among neo-challenged and re-challenged conditions. F. Treg frequency among neo-challenged and re-challenged conditions. G. Study of immune checkpoint expression in Tregs among neo-challenged and re-challenged conditions. H. Study of CD49d^+^CD11a^+^ tumor specific CD4^+^ and CD8^+^ T cells in neo-challenged and re-challenged conditions, \**p*.val <0.05; \*\**p*.val<0.01, \*\*\**p*.val<0.001.

Next, we investigated activation marker expression on CD4^+^ and CD8^+^ T cells (Fig8.E, F) from LN. In CD4^+^ T cells no modification in the expression of CD40L, IL-2Rβ or 4-1BB was observed. However, the expression of ICOS in re-challenged mice after anti-HVEM18-10 (7.017% ± 1.22) or anti-CTLA-4 (8.51% ± 1.82) treatment increased compared to controls (3.43% ± 0.32). Similarly, CD69^+^CD4^+^ T cells were more abundant in re-challenged mice (anti-HVEM18-10: 15.56% ± 1.57; anti-CTLA-4: 19.4% ± 1.93) compared to controls (11.64% ± 1.24). Noteworthy, CD69 overexpression was greater in anti-CTLA4 re-challenged mice compared to anti-HVEM re-challenged mice (Fig8.E). In CD8^+^ T cells, an over expression of IL2-Rβ in re-challenged mice (anti-HVEM18-10: 30.16% ± 1.97; anti-CTLA-4: 25.2% ± 3.94) compared to controls (7.13% ± 0.91) was observed. Here, IL2-Rβ expression was greater in anti-HVEM18-10 re-challenged mice compared to anti-CTLA-4 re-challenged mice (Fig8.F). This suggests a better response to IL-2 and consequently more CD8^+^ T cell activation. Noteworthy, we did not notice any difference in the expression of CD69 on CD8^+^ T cells (Fig8.F). Next, the Treg population within LNs after re-challenge was investigated. Treg frequency increased after re-challenge in both anti-HVEM18-10 (7.91% ± 2.29) and anti-CTLA-4 (14.4% ± 2.47) mice compared to anti-HVEM18-10 neo-challenged group (2.46% ± 0.36) (Fig8.G). Last, a decrease in the expression of CD69 in Treg was observed, suggesting less activation or recruitment of peripheral blood Treg (Fig8.H). Taken together, these results show that re-challenged mice rejected secondary tumor inoculation thanks to a strong T cell memory response linked to an increased expression of IL-2Rβ and a recruitment of tumor responsive CD8^+^ and CD4^+^ T cells.

Recently, the CD11a^+^CD49d^+^ CD4^+^ and CD8^+^ T cells were reported as tumor antigen responsive T cells, which participate to anti-tumor response (23). Therefore, we gated CD11a^+^CD49d^+^ CD4^+^ and CD8^+^ T cells (Fig8.D) and showed that tumor antigen responsive CD4^+^ T cells increased upon re-challenge with anti-HVEM18-10 (9.51% ± 0.38) and anti-CTLA-4 (10.71% ± 1.9) mice (Fig8.D left) compared to controls (1.72% ± 0.24). Interestingly, tumor antigen responsive CD8^+^T cells significantly increased after re-challenge with anti-HVEM18-10 (1.72% ± 0.24) only (Fig8.D right). Taken together, our results show that the anti-tumoral memory response that allowed tumor rejection relies on enrichment in CD4^+^ and CD8^+^ memory subpopulations, increased IL-2Rβ and activation markers and more importantly on tumor antigen specific T cell subsets, which most likely allow secondary tumor rejection.

## Discussion

HVEM is a molecular switch as its effect depends on the ligand involved, largely expressed in lung and colorectal tumors. Here, we showed that our anti-HVEM18-10 antagonist mAb potentiate *in vitro* T cell proliferation and activation by blocking preferentially the interaction with the inhibitory ligands of HVEM. Then, we developed an innovative mouse model expressing both human BTLA and human HVEM. Anti-HVEM18-10 injection in mice induced the development of a marked T cell-memory phenotype and Tumor antigen specific (CD49d^+^CD11a^+^) T cells contributing in HVEM^+^ a tumor reduction or rejection. Therefore, HVEM targeting is a great addition to the currently available arsenal of immunotherapies.

To assess the clinical relevance of HVEM targeting in lung and colorectal tumors, we screened HVEM expression public transcriptomic databases. We found that HVEM was largely expressed in these cancers and did not correlate with PD-L1 expression. Surprisingly, the literature reporting HVEM expression in lung and colorectal cancers remains sparse. To our knowledge, HVEM expression on NSCLC biopsies were studied once by IHC showing that HVEM was overexpressed in patients with advanced disease and was negatively correlated to PD-L1 expression (14). In colorectal cancer, two studies by IHC reported a correlation between HVEM overexpression and advanced disease and poorer prognosis (8,24). Nevertheless, a more detailed multi-parametric study would improve our knowledge of HVEM expression and regulation by the different immune and non-immune actors of colon and lung TME, similarly to Steele *et al*. study in pancreatic cancer (25). This would participate to the improvement of tumor type stratification and define who could benefit from HVEM18-10 mAb treatment.

In our *in vitro* pre-clinical settings, we showed that the anti-HVEM18-10 mAb increased primary human αβ-T cells activity alone (CIS-activity) or in presence of HVEM-expressing lung or colorectal cancer cells *in vitro* (TRANS-activity). Thus, we observed an immune reactivation similarly to that triggered by anti-PD-1 or anti-CTLA-4 treatment, favoring the idea that HVEM18-10 mAb avoids HVEM interaction with its inhibitory ligands. The same observation was made with anti-HVEM18-10 which allowed γδ-T cells activation, especially against HVEM^+^ lymphoma cells (15). In the light of these observations, the use of HVEM18-10 mAb to potentiate anti-tumor T cell responses in hematological cancer as well as in solid cancers is of great interest.

Interestingly, anti-HVEM18-10 synergized with anti-PD-L1 effects and maximized T cells responses when tumor cells expressed both targets. Therefore, anti-HVEM18-10 could be used in combination with other immune checkpoint inhibitor therapies and reinforce the still growing arsenal of ICI combinations in on going trials, which includes among others anti-CTLA4/antiPD-L1 (26) anti-TIGIT/anti-PD-L1 (27), and anti-BTN3A1/anti-PD-L1 (28). Nevertheless, we showed that anti-HVEM18-10 alone is still sufficient to trigger efficient T cells responses in PD-L1^-^ conditions. This is of particular interest in cancers such as colorectal cancer, where PDL1 is rarely expressed and the clinical benefits of antiPD1/PDL1 immunotherapies restricted to a small subset of patients (3)(4). Thus, anti-HVEM18-10 may increase the target patient population independently from their PDL1 status.

The improvement of *in vitro* anti-tumor T cell function led us to evaluate the efficacy of anti-HVEM18-10 in animal settings. Recently, anti-HVEM IT was investigated in a prostate cancer mouse model (29) where Aubert *et al*. showed that anti-HVEM18-10 reduced the growth of a HVEM^+^tumors. Their mouse model, a NOD.SCID.gc-null mice reconstituted with human T cells, demonstrated that anti-HVEM18-10 efficacy was mainly mediated by the infiltration of CD8^+^ T cells. Moreover, anti-HVEM18-10 increased TIL number, proliferation and enriched pathways associated to lymphocyte activation, such as TNF mediated signaling pathway or TCR complex. Finally, genes involved in immunosuppressive pathways such as *BTLA, TIGIT, LAG3, TIM3*,, were downregulated in the anti-HVEM18-10 treated group (29). In our study, we generated innovative immunocompetent mouse models expressing extracellular domains of human BTLA or both human BTLA and human HVEM, while the intracellular domains and signaling remained murine. This, allowed pre-clinical experiments settings using human anti-HVEM18-10 in immunocompetent mice bypassing the use of human T cell reconstitution in nude mice, which strengthened observations. We demonstrated that anti-HVEM18-10 strongly delays human HVEM^+^ tumor growth and led to tumor eradication in 20% of mice. TIL phenotype showed a decrease in Treg infiltration and exhausted CD8^+^ T cells, and in increase of EM CD4^+^ T cells, expanding results from Aubert *et al*. in prostate tumors (29) to colon cancer. Therefore, we highlighted a common way of activating lymphocytes in different solid tumor models. In addition, anti-CTLA4 treatment showed similar phenotypic modifications, which was already reported in the literature (21,22). Altogether, these data suggest that the local immune response is strengthened by anti-HVEM18-10 IT thanks to a suppression of intra-tumoral exhausted CD8^+^ T cell and Treg populations, which slowed tumor growth and even eradicated the tumor. Finally, our new mouse models allowed the study of therapeutic mAb directly in an immunocompetent and dynamic environment. To date only one recent study described a KI mouse model expressing Human PD-1 and CTLA-4 molecules (30). The number of these syngeneic mouse models will increase in the future because they are closer to the clinic as they allow the use of anti-human target mAbs and allow a precise investigation of mAb effects.

In our settings, around 20% of the mice completely rejected the tumor enabling the study of memory response induced by the re-challenge. Indeed, tumor rejecting mice were re-challenged in the contralateral flank with a larger amount of tumor cells and still rejected the tumor. Within tumor draining lymph nodes, we found a specific T cell memory composition, marked by an increase in IL-2Rβ^+^ CM CD8^+^ T cells and EM CD4^+^ T cells in comparison to neo-challenged mice. More importantly, draining lymph node sheltered a population of CD49d^+^CD11a^+^ CD8^+^ and CD4^+^ lymphocytes, which were recently described as tumor antigen specific T cells (31). after anti-HVEM18-10 treatment. Therefore, anti-HVEM18-10 treatment leads to a systemic, tumor antigen specific, T cell memory response allowing the mice to control distal secondary tumor implantation. Similar to our model, CD8^+^ T cells showed a marked memory phenotype and response better to secondary stimuli under anti-CTLA4 or anti-PD1 treatment in other studies (32–35).

## Conclusion

HVEM appears to be a very promising IT target for oncologic and hematologic malignancies. Anti-HVEM18-10 mAb treatment demonstrated that anti-tumor immune response was strengthened, delays tumor growth or eradicate tumors and induces a memory immune response in a cutting-edge pre-clinical mouse model. Altogether, these results highlight the interest of anti-HVEM IT alone or in combination with another IT to further enhance anti-tumor immunity.

## Acknowledgements

CIPHE is supported by the Investissement d’Avenir program PHENOMIN (ANR-10-INBS-07) This study was supported by the RHU PIONEER (ANR-17-RHUS-0007), France 2030 LG and CD are supported by the RHU PIONEER (ANR-17-RHUS-0007), France 2030

