## Supplemental figures for "Anti-HVEM mAb therapy improves antitumoral immunity both *in vitro* and *in vivo*, in a novel transgenic mouse model expressing human HVEM and BTLA molecules challenged with HVEM expressing tumors"

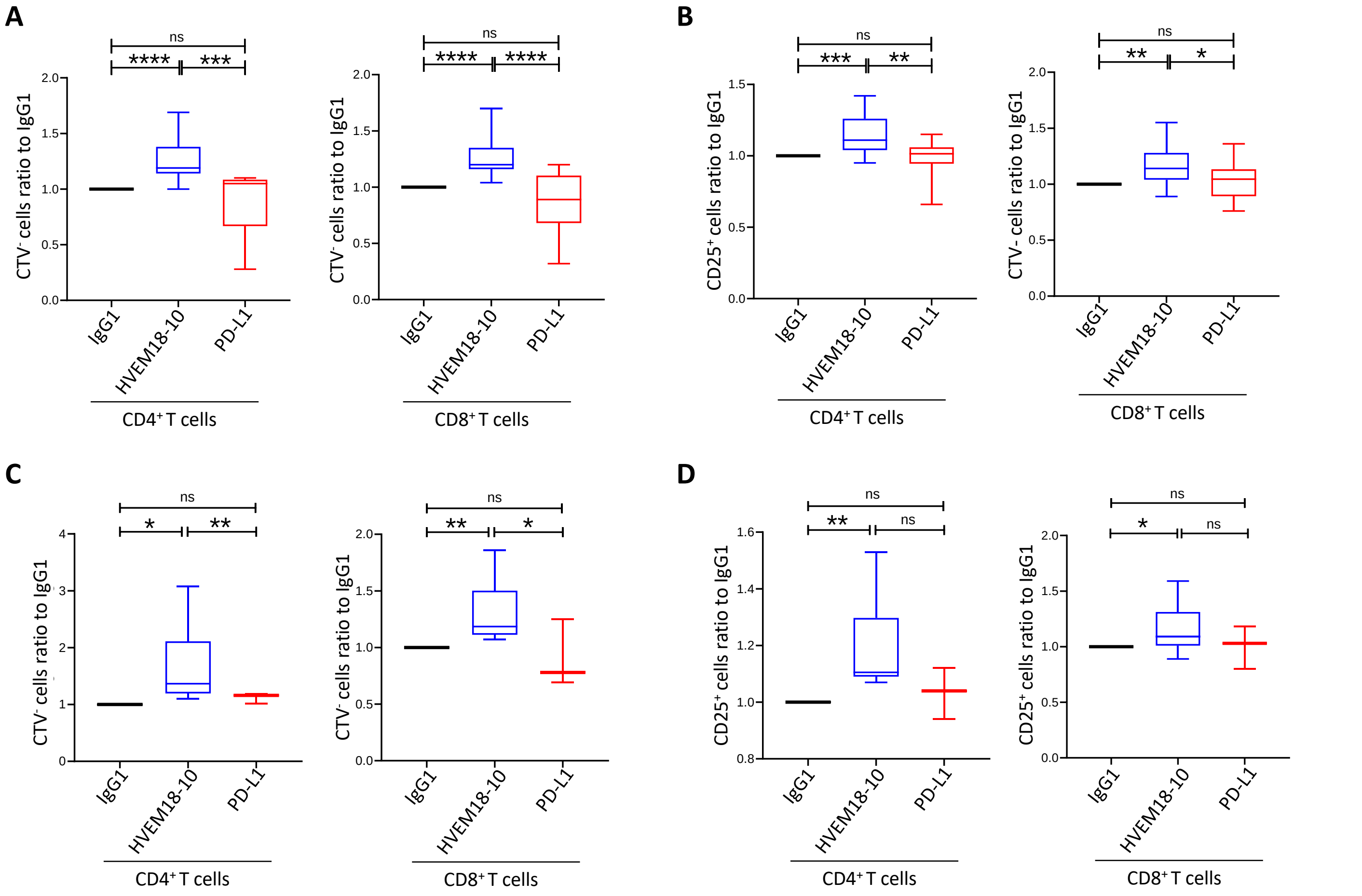


**Supp.Figure.1 anti-HVEM18-10 enhances T cells response against PD-L1 negative lung and colorectal cancer cell line**

PD-L1 negative lung cancer NCIH2405 (A-B) or colorectal cancerHT29 (C-D) cell lines were seeded 24hours before the experiment. Then, cultured PBMC from healthy donors for 72h with OKT3 stimulation and treated or not (igG1) with anti-HVEM 18-10 antibody (blue bars/lines) or anti-PD-L1 (red bars). (A,C) proliferation profile of T cells by Cell TraceViolet staining (A for NCIH2405 and C HT29) and CD25 expression (B for NCIH2405 and D for HT29). Bar plots are the Mean ± SEM of different healthy donors samples, (A-B n=15; C-D : HVEM18-10 condition n=8 ; PD-L1 condition n =3). * $p < 0.05$, ** $p < 0.01$, *** $p < 0.005$, **** $p < 0.001$(Student's t-test)

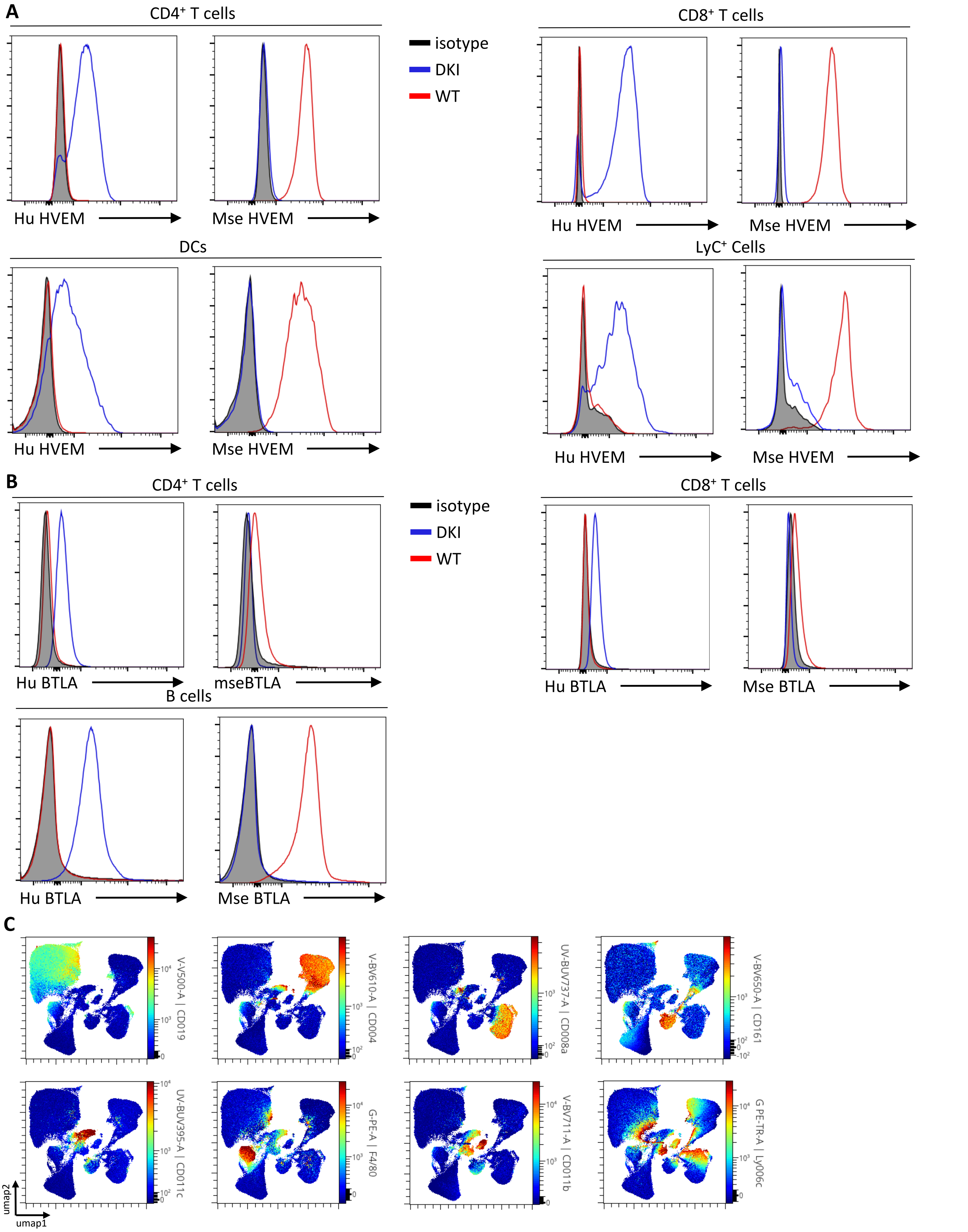


**Supp.Figure 2 (associated to Fig5): hBTLA$^{+/+}$ and hBTLA$^{+/+}$hHVEM$^{+/+}$ Mice show similar immune cell phenotypes.** A. Human or mouse HVEM expression was assessed on CD4+ , CD8+ T cells, DCs (CD11c+MHC class II+) and LyC+ cells from DKI (Blue),WT mice (red) or isotypic Ig control (black) B. Human or mouse HVEM expression on CD4+,CD8+ T and B cells cells from DKI (Blue),WT mice (red) or isotypic Ig control (black) C.Individual marker expression for the gating of each individual immune cell population.

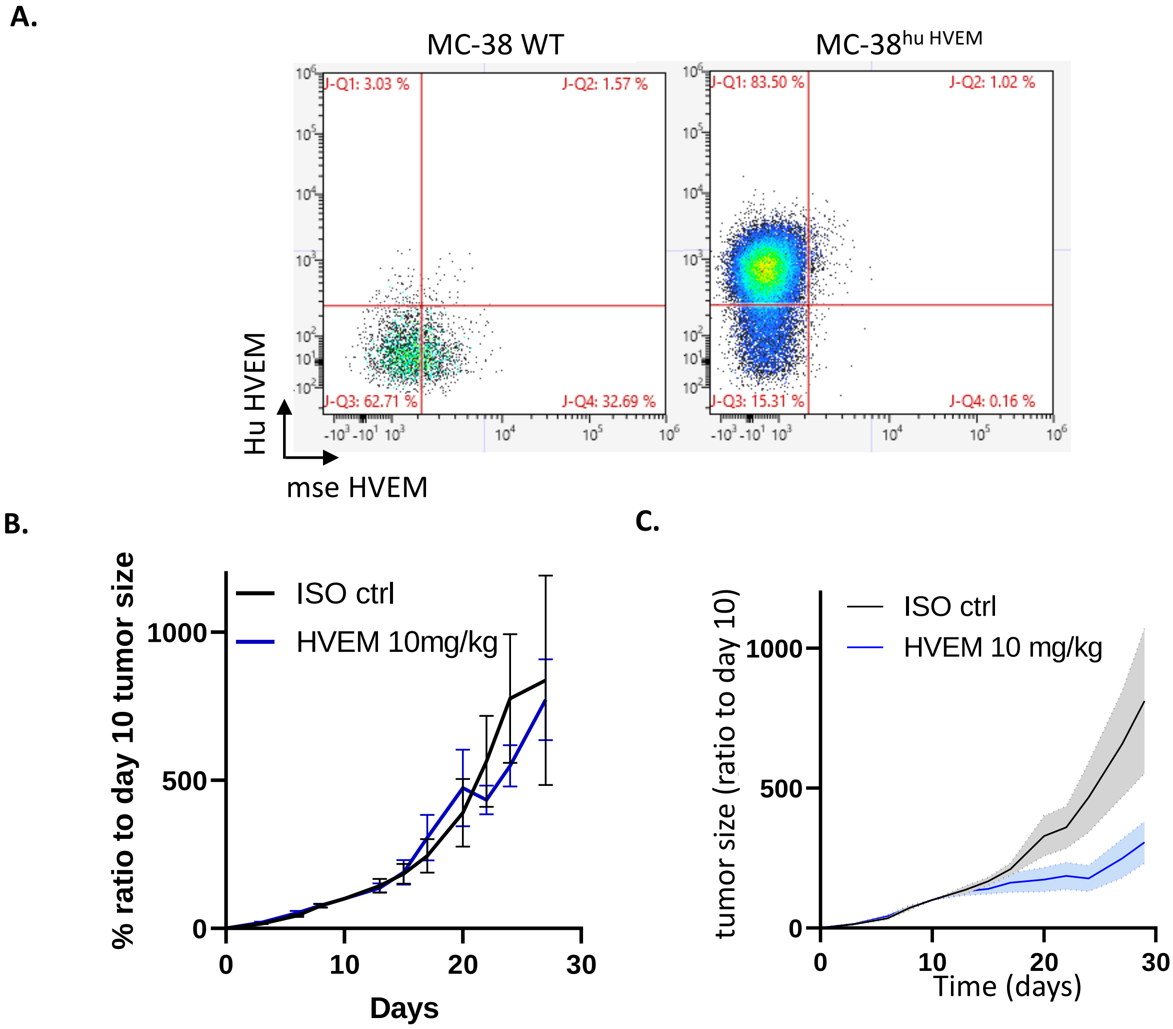


Supp.Figure 3 blocking Trans-BTLA-HVEM binding *in vivo* is sufficient to decrease solid tumor growth
A.MC38-WT or MC-38 1E10 clone expression of Human or mouse HVEM in cytometry B.Measure of tumor growth showed a ratio to size of tumor at randomization day. Colorectal cancer cells MC-38hu HVEM were injected ($0.5 \times 10^6$ ) at day 0, and Iso ct (black) or anti-HVEM 18-10 antibody 10 mg/kg (blue).

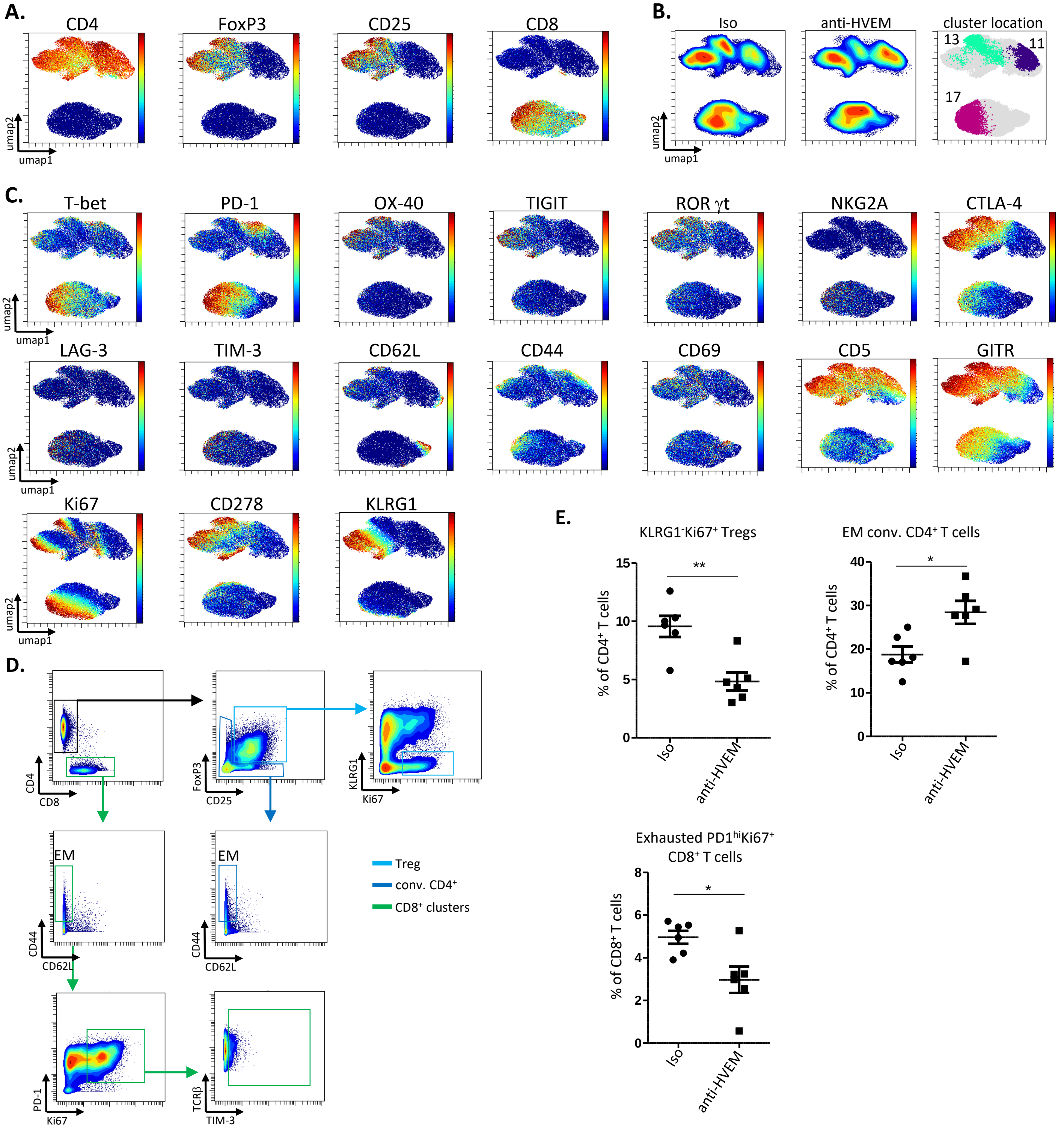


**Supp Figure 4 (associated to Fig7) : Individual marker expression, significantly modulated cluster location and manual gating validation of clusters 11, 13 and 17.**

**A.** Main gating marker UMAP for the identification of Tregs, conventional $CD4^+$ T cells and $CD8^+$ T cells. **B.** Density UMAP and significantly altered cluster location. **C.** Phenotype and function marker UMAPs used to generate the heat map in Figure 8. **D.** Strategy allowing the gating of PhenoGraph clusters 11 (EM conv. $CD4^+$ T cells), 13 ($KLRG1^-Ki67^+$ Tregs) and 17 (exhausted PD-$1^{hi}Ki67^+CD8^+$ T cells) for manual validation. **E.** Statistical validation of manually gated clusters (from D). **p*.val <0.05; ***p*.val<0.01.

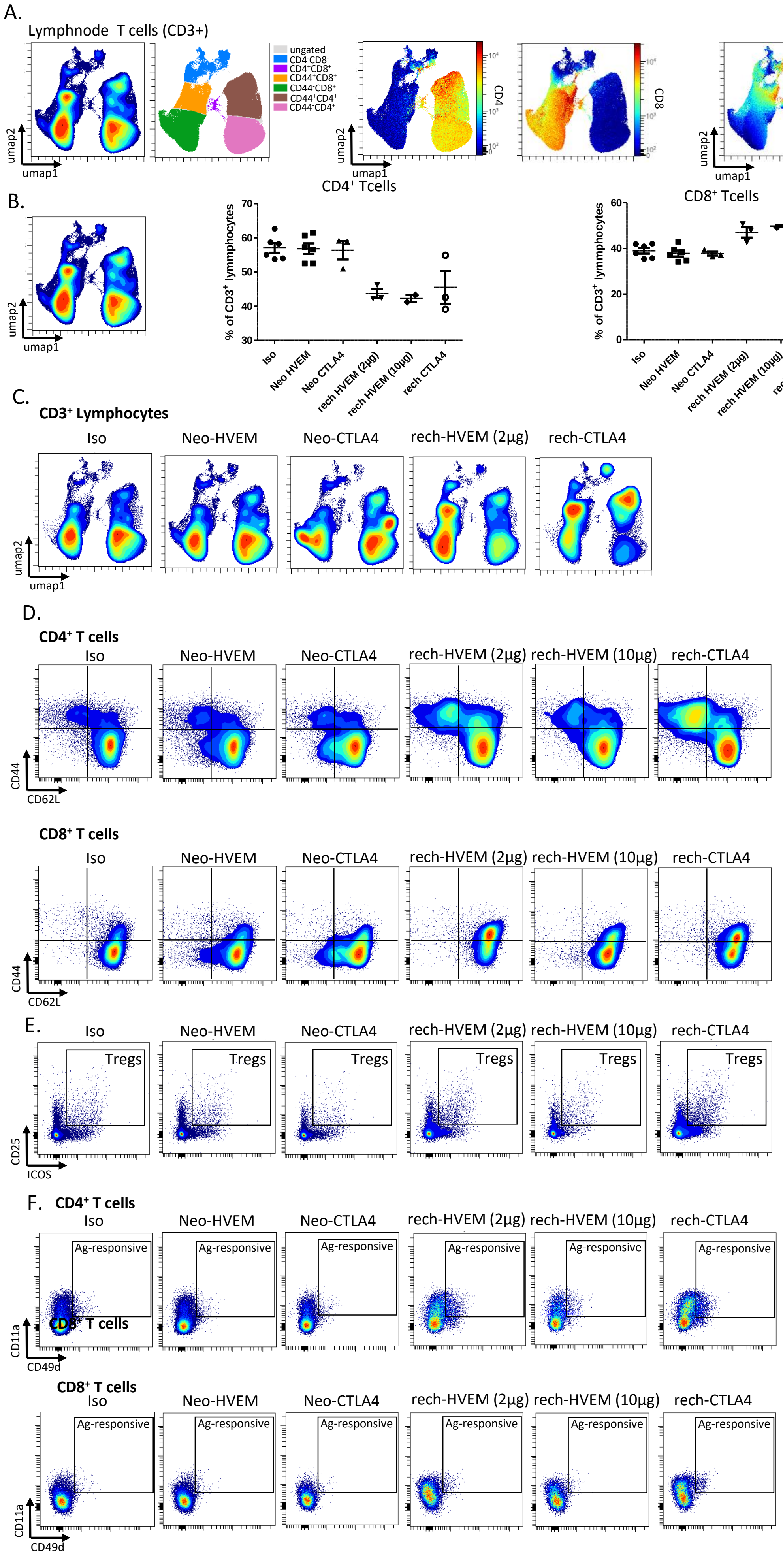


**Supp Figure 5 (associated to Fig8) : re-challenged mice exhibit specific T cell phenotypes**

**A.** T cell clusters in mice lymphnodes were generated using flow cytometry data, T cell annotation was performed using CD4, CD8, CD25 and CD44 expression. **B.** Frequency analysis of $CD4^+$ and $CD8^+$ T cell in neo- and re-challenged conditions. **C.** Density analysis of T cells in isotype, anti-HVEM18-10 and anti-CTLA-4 neo- and re-challenged mice. **D.** Manual gating for EM, CM and naive $CD4^+$ and $CD8^+$ T cells in each condition. **E.** Manual gating of Tregs in each condition. **F.** Manual gating of antigen responsive ($CD49d^+CD11a^+$) $CD4^+$ and $CD8^+$ T cells.
